# A family of endonucleases that block nanotube-mediated plasmid dissemination

**DOI:** 10.1101/2025.10.26.684598

**Authors:** Venkadesaperumal Gopu, Saurabh Bhattacharya, Michal Bejerano-Sagie, Mei Zhuang, Yuval Nevo, Bushra Shraiteh, Miriam Ravins, Manas Kumar Guria, Tamar Kahan, Ilan Rosenshine, Sigal Ben-Yehuda

## Abstract

Small non-conjugative plasmids constitute a substantial portion of the bacterial mobile genome, driving the dissemination of beneficial genes, with their transfer primarily attributed to transformation, transduction, or co-mobilization with conjugative elements^1–3^. Here we explore an understudied plasmid spread route among bacteria, mediated by intercellular membranous nanotube conduits^4^. We reveal that, unlike traditional donor-to-recipient delivery, **N**anotube-dependent **P**lasmid **ex**change (NPex) operates bidirectionally, enabling both plasmid donation and, to a lesser extent, plasmid acquisition. By identifying a *Bacillus subtilis* natural isolate deficient in NPex, we discovered a prophage-encoded factor, YokF, that blocks plasmid transmission, chiefly acting within the donor cell to inhibit plasmid donation. YokF is a nuclease that localizes to the membrane, where it interacts with a nanotube component to selectively impede plasmid transfer through degradation. Importantly, YokF homologs from various bacterial species were found to exhibit anti-NPex activity. Given their prevalence in Gram-positive bacteria, we propose that YokF homologs represent a conserved family of NPex gatekeepers that restrict plasmid flow within bacterial communities.

## Introduction

Plasmids play a pivotal role in bacterial ecology and evolution by facilitating the horizontal transfer of accessory genes, including those encoding antibiotic resistance, metabolic traits, and virulence factors, thereby significantly contributing to bacterial adaptability, resilience, and pathogenesis [e.g.,^5–11^]. The exchange of large plasmids among bacteria is primarily driven by conjugation, where a dedicated plasmid-encoded secretion system forms an intercellular physical bridge, providing a path for donor-to-recipient plasmid delivery [e.g.,^12–14^]. However, the bacterial plasmidome exhibits remarkable diversity, and many small-size non-conjugative plasmids, lacking dedicated transfer machinery, are widely prevalent in nature [e.g.,^15–18^]. The spreading of these small plasmids into new hosts is thought to occur via transformation, transduction, or co-mobilization with conjugative elements cohabiting with the same host [e.g.,^1–3^].

In recent years, it has become apparent that plasmids can spread among bacterial communities through mechanisms exceeding traditional routes. DNA can be packaged and transported over long distances by ubiquitously produced membrane vesicles [e.g.,^19–21^]. Further, intercellular membranous conduits, termed nanotubes, were found to facilitate the exchange of non-conjugative plasmids, along with additional cargo including small molecules, nutrients, and proteins^4,22–27^. Nanotube-like structures are widespread in bacteria, predominantly when grown on solid surfaces or in biofilm assemblies, and can also be found in eubacteria, archaebacteria, and eukaryotes [e.g.,^23,28–36^]. Recently, nanotube networks connecting marine cyanobacteria species have been described^37^, highlighting their potential universal impact on bacterial communities in varied niches. Here, we describe a family of proteins that act to prevent nanotube-dependent intercellular plasmid transmission.

## Results

### Nanotube-dependent transfer of small non-conjugative plasmids

Natural *Bacilli* species carry a plethora of small non-conjugative plasmids^17,38–41^, we thus examined the impact of intercellular nanotubes on their spread from donor to recipient bacteria. In accord, *B. subtilis* donor cells (PY79), harboring plasmids of different origins (3.4, 5.6, and 6.6 kb), were mixed with plasmid-free recipient bacteria, and plasmid transfer efficiency was estimated at 4.8×10^−6^ - 6.2×10^−7^, indicating the recipient fraction obtaining a plasmid (termed trans-recipients) (Fig. 1A-1B; Table S1). Similar results were obtained using the naturally abundant *B. subtilis* plasmid pTA1015 (9.2 kb)^38,40^ (Fig. 1A-1B). Nanotube biogenesis was found to depend on the conserved flagellar export apparatus, comprising FlhA, FlhB, FliP, FliQ, and FliR, collectively designated CORE. This complex localizes to the base of the nanotube, alternatively functioning as a platform for flagella or nanotube formation^26,42,43^. We thus tested plasmid exchange between *CORE* deficient mutants. Plasmid exchange was strictly CORE-dependent and independent of motility, the competence machinery, transduction mechanisms, or membrane vesicles^26,44,45^ (Fig. 1A-1B; Fig. S1A-S1B). Hence, we termed this mode of plasmid delivery “**N**anotube-dependent **P**lasmid **ex**change (NPex)”. Nevertheless, while using a large plasmid pLS20 (∼65 kb), lacking a functional conjugation system (Δ*virB4*), no plasmid exchange was detected (Fig. S1C-S1D), suggesting size constraints for NPex. Overall, NPex represents a significant path for the spread of small plasmids among *B. subtilis* cells.

**Fig. 1.**
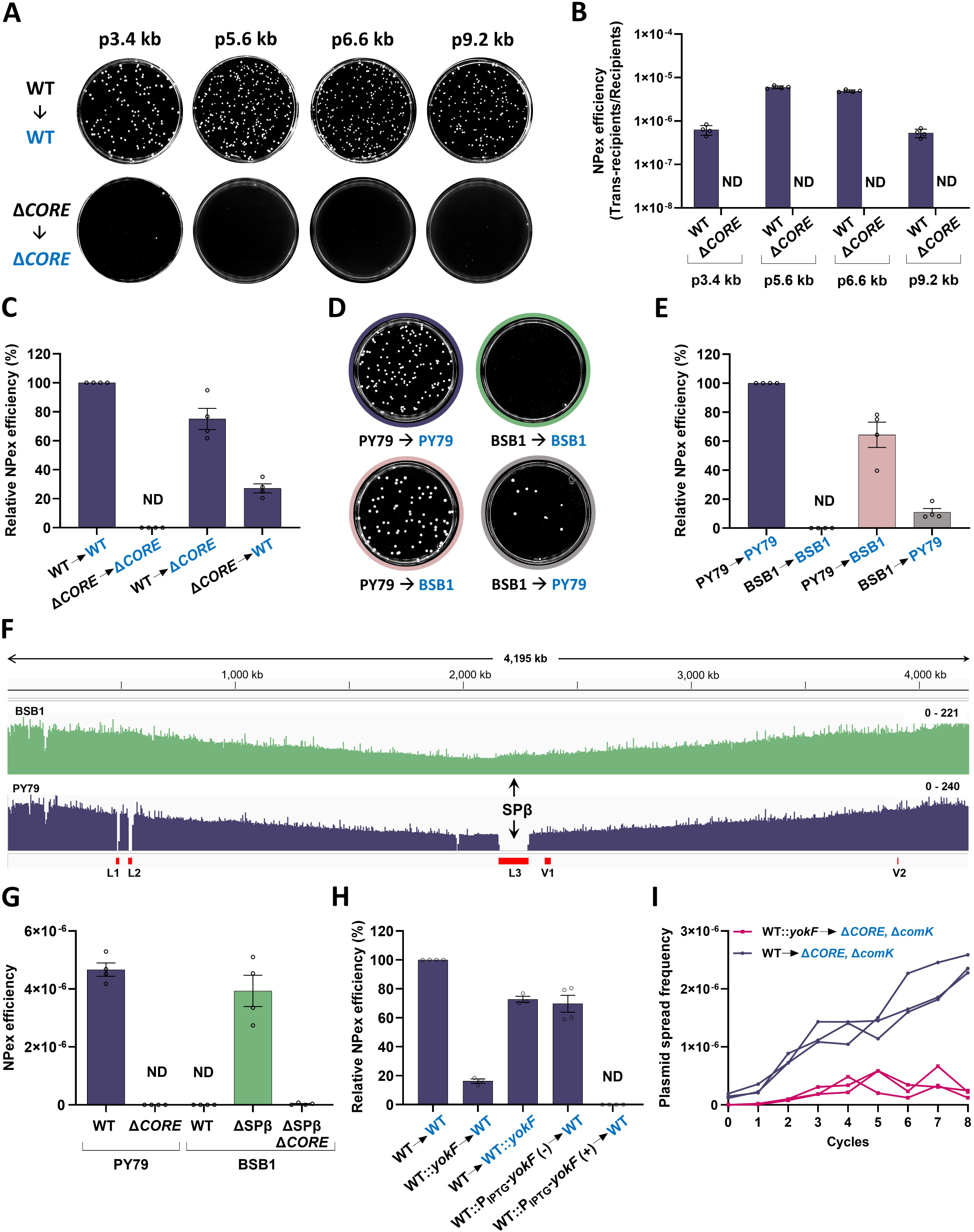
Nanotube-dependent plasmid exchange (NPex) and its inhibition by YokF. **(A)** <u>NPex is *CORE*-dependent</u>: Upper panels: WT (PY79) donor strains harboring small non-conjugative plasmids [p3.4 kb (pNZ8048/*cat*), p5.6 kb (pCPP31/*cat*, *neo*), p6.6 kb (pHB201/*cat*, *erm*), and pTA1015-*cat* (p9.2 kb/*cat*)], were mixed in 1:1 ratio with WT (PY79) recipients harboring either kanamycin (*sacA*::*kan*) or erythromycin (*sacA*::*mls*) resistance genes. The mixtures were incubated for 4 hours without selection, and equal volumes of cells were then spread over plates selective for trans-recipients. Lower panels: A similar experiment was conducted using equivalent Δ*CORE* donor and recipient strains. A detailed description of the used donor (black) and recipient (blue) pairs, and antibiotic selections employed, are listed in Table S1. Shown are representative images of NPex selective plates. **(B)** Quantitative analysis of NPex efficiencies between donor and recipient pairs shown in (A). NPex efficiency=Trans-recipients CFU/Total recipients CFU. Data are presented as mean values and SEM, based on 4 independent experiments. ND – Not Detected. **(C)** <u>NPex directionality</u>: WT (PY79) (GV478) or isogenic Δ*CORE* (GV491: Δ*fliO-flhA*::*tet,* P*_fla/che_*-*flhF*) donor strains carrying p6.6 kb (pHB201/*cat*, *erm*), were mixed in 1:1 ratio with WT (SB513: *amyE*::P_IPTG_-*gfp*-*kan*) or isogenic Δ*CORE* (SH13: Δ*fliO-flhA*::*tet,* P*_fla/che_-flhF*, *amyE*::P_IPTG_-*gfp*-*kan*) recipients, harboring kanamycin resistance gene. The used donor (black) and recipient (blue) pairs are indicated. The mixtures were incubated for 4 hours without selection, and equal volumes of cells were then spread over plates containing chloramphenicol, erythromycin, and kanamycin to select for trans-recipients. NPex efficiency was determined as in (B). Shown is NPex efficiency (%) relative to WT → WT pair. Data is presented as mean values and SEM, based on 4 independent experiments. ND – Not Detected. **(D)** <u>BSB1 is NPex deficient</u>: PY79 (GV478) or BSB1 (GV345) donor strains carrying p6.6 kb (pHB201/*cat*, *erm*), were mixed in 1:1 ratio with PY79 (SB513: *amyE*::P_IPTG_-*gfp*-*kan*) or BSB1 (GV372: *amyE*::P_IPTG_-*gfp*-*kan*) recipients, harboring kanamycin resistance gene. The used donor (black) and recipient (blue) pairs are indicated. The mixtures were incubated for 4 hours without selection, and equal volumes of cells were then spread over plates containing chloramphenicol, erythromycin, and kanamycin to select for trans-recipients. Shown are representative images of NPex selective plates. **(E)** Quantitative analysis of NPex efficiencies between donor and recipient pairs shown in (D). NPex efficiency was determined as in (B). Shown is NPex efficiency (%) relative to PY79 → PY79 pair. Data is presented as mean values and SEM, based on 4 independent experiments. ND – Not Detected. **(F)** <u>A comparison between BSB1 and PY79 genomes</u>: BSB1 (green) and PY79 (purple) strains were subjected to whole genome sequence analysis. The Y-axis represents the number of sequence reads, whereas the X-axis shows the chromosomal position. Variable regions between the two genomes (V1 and V2), and regions lacking from the PY79 genome (L1-L3) are indicated (red). The location of SPβ (L3) is specified by arrows. The scheme was generated using Integrative Genomics Viewer (IGV). **(G)** <u>SPβ prophage inhibits NPex</u>: Donor strains: WT (PY79), isogenic Δ*CORE*, WT (BSB1), BSB1 lacking SPβ, and BSB1 lacking both SPβ and *CORE*, all carrying the p6.6 kb (pHB201/*cat*, *erm*), were mixed in a 1:1 ratio with kanamycin resistance recipient strains, harboring equivalent genotypes, but lacking a plasmid. The mixtures were incubated for 4 hours without selection, and equal volumes of cells were then spread over plates selective for trans-recipients. A detailed description of the used donor and recipient pairs, and antibiotic selections employed, are listed in Table S1. NPex efficiency was calculated as in (B). Data is presented as mean values and SEM, based on 4 independent experiments. ND – Not Detected. **(H)** <u>YokF inhibits NPex</u>: Donor strains: WT (PY79) (GV478), PY79 expressing YokF from its native promoter WT::*yokF* (GV469: *sacA*::P*_yokF_*-*yokF*_BSB1_-*spec*), or from an IPTG-inducible promoter WT::P_IPTG_-*yokF* (GV480: *amyE*::P_IPTG_*-yokF*_BSB1_-*spec*), all carrying p6.6 kb (pHB201/*cat*, *erm*), were mixed in a 1:1 ratio with kanamycin resistance recipient strains: WT (SB513: *amyE*::P_IPTG_-*gfp*-*kan*), or WT::*yokF* (GV468: *sacA*::P*_yokF_*-*yokF*_BSB1_-*spec*, *amyE*::P_IPTG_*-gfp-kan*). The used donor (black) and recipient (blue) pairs are indicated. Mixtures were incubated for 4 hours without selection in the presence (+) or absence (-) of IPTG. Equal volumes of cells were then spread over plates containing chloramphenicol, erythromycin, and kanamycin to select for trans-recipients. NPex efficiency was determined as in (B). Shown is NPex efficiency (%) relative to WT → WT pair. Data is presented as mean values and SEM, based on at least 3 independent experiments. **(I)** <u>YokF limits plasmid dissemination</u>: Donor strains (black), WT (PY79) (GV478) or PY79 expressing YokF from its native promoter WT::*yokF* (GV469: *sacA*::P*_yokF_*-*yokF*_BSB1_), carrying p6.6 kb (pHB201/*cat*, *erm*) were mixed in 1:1 ratio with kanamycin resistance recipient (blue) Δ*CORE,* Δ*comK* (GV645: Δ*fliO-flhA::tet,* P*_fla/che_-flhF*, Δ*comK*::*kan*, *sacA*::P*_veg_*-*mCherry*). The mixtures were incubated for 4 hours with sublethal concentration of lincomycin. Equal volumes of cells were then spread over plates containing chloramphenicol, erythromycin, and kanamycin to select for trans-recipients, and on plates containing only kanamycin to select for the total recipient population (Cycle 0). Subsequently, the recipient population was collected and introduced to new donor cells and plasmid spread efficiency was followed over 8 consecutive cycles. The graph illustrates plasmid spread frequency, determined as the ratio of trans-recipients (CFU)/total recipient population (CFU) for each cycle. Data is presented as absolute values, based on 3 independent experiments.

To gain insight into NPex directionality, we crossed wild-type (WT) donor with nanotube impaired Δ*CORE* recipient, and assayed for NPex, using p6.6 kb (pHB201) as our standard NPex marker. The mixture yielded a slight reduction in NPex events to approximately 75±12% of those observed for the WT cross (Fig. 1C). Intriguingly, when crossing Δ*CORE* donor with a WT recipient, approximately 27±5% of NPex events could still be detected (Fig. 1C). These results signify that, unlike conjugation, NPex operates bidirectionally, as both donor and recipient are capable of projecting nanotubes. The primary route is executed by the nanotube-producing plasmid-containing donor, whereas the minor route is performed through active plasmid acquisition by the nanotube-producing recipient.

### Prophage-encoded YokF inhibits NPex

When assessing NPex manifestation across a range of *B. subtilis* strains, we found the commonly used BSB1 strain to be completely NPex impaired (Fig. 1D-1E)^46,47^. Curiously, crossing PY79 strain as a donor with BSB1 recipient largely rescued the observed deficiency (64±15%) (Fig. 1D-1E). When BSB1 served as a donor to PY79 recipient, a low exchange rate was detected (11±4%) (Fig. 1D-1E), implying that BSB1 phenocopies the PY79 Δ*CORE* mutant (Fig. 1C). To uncover potential mutations impairing NPex execution by BSB1, we compared PY79 and BSB1 genotypes following whole genome sequencing. Two regions, *panB-hepT* (V1) and *sacA-ywcI* (V2), contained multiple single nucleotide polymorphisms (Fig. 1F). Nonetheless, swapping these variable regions between the strains neither enabled BSB1, nor interfered with PY79, NPex capability (Fig. S1E; Table S1). This led us to consider that rather than lacking a critical NPex function, BSB1 might harbor a factor actively impeding NPex. Notably, the BSB1 contains three genomic regions, absent from PY79 (L1-L3): a 16 kb region, *lrpC*-*mntH*, (L1), an ICE1 conjugative element (L2), and SPβ prophage (L3)^48,49^ (Fig. 1F). Remarkably, deleting the SPβ prophage from the BSB1 genome (ΔSPβ), but not the L1 or L2 regions, unveiled BSB1 NPex proficiency to a level similar to that of PY79 (Fig. 1G; Fig. S1F; Table S1). Importantly, this regained ability was *CORE*-dependent (Fig. 1G; Table S1), insinuating that SPβ harbors gene(s) that actively inhibit NPex. Notably, this finding resolves the discrepancy, in which BSB1 failed to exhibit NPex^47^, most likely due to the masking effect of the SPβ prophage.

Deletion analysis of the prophage genome pinpointed *yokF* as the key contributor to the NPex halting phenotype (Fig. S1G-S1H; Table S1). Ectopic expression of *yokF* in the PY79 donor, under its native promoter, effectively blocked approximately 83±2% of NPex events, while its expression in the recipient mildly (27±3%) influenced the process, similar to that of Δ*CORE* phenotype (Fig. 1C and 1H). Furthermore, donor-produced YokF from an inducible promoter entirely ceased NPex (Fig. 1H). Of note, YokF expression had no significant influence on competence, phage sensitivity, or plasmid stability (Fig. S1I-S1K). Hence, YokF expression in donor bacteria is sufficient to block the NPex process, in a manner that is independent of any additional SPβ factors. Surprisingly, YokF did not seem to interfere with nanotube formation nor to perturb nanotube-mediated protein exchange (Fig. S2A-S2B), suggesting a specific effect on plasmid cargo.

To determine YokF impact on plasmid dissemination, we examined the kinetics of plasmid spread mediated by PY79 donor strains either lacking or harboring *yokF* produced from its natural promoter. As a recipient, we employed an NPex and competence deficient mutant (Δ*CORE,* Δ*comK*), and imposed mild sub-lethal antibiotic pressure to enhance delivery events. Following multiple crossing cycles, the proportion of recipient bacteria carrying the plasmid significantly increased in the absence of YokF, while remaining low in its presence (Fig. 1I). *In toto*, YokF operates to block NPex, profoundly reducing plasmid spread dynamics within bacterial communities.

### YokF displays nuclease activity crucial for NPex inhibition

To gain insight into YokF functionality, we conducted bioinformatic analysis combined with AlphaFold and HHpred^50,51^ to predict the protein structure. YokF contains a putative N-terminal signal peptide (SP) (1-20 aa), indicating probable membrane or surface localization, and a consequent central globular thermo-nuclease domain (TNase) (66-199 aa) capable of catalyzing DNA hydrolysis (Fig. 2A). The TNase was succeeded by an unstructured region (216-255 aa) proceeding a globular C-terminal domain (256-296 aa) (Fig. 2A). The latter appears to dock over and potentially modulate the TNase function (Fig. 2A), hence we refer to it as the TNase modulator (TNM) domain.

**Fig. 2.**
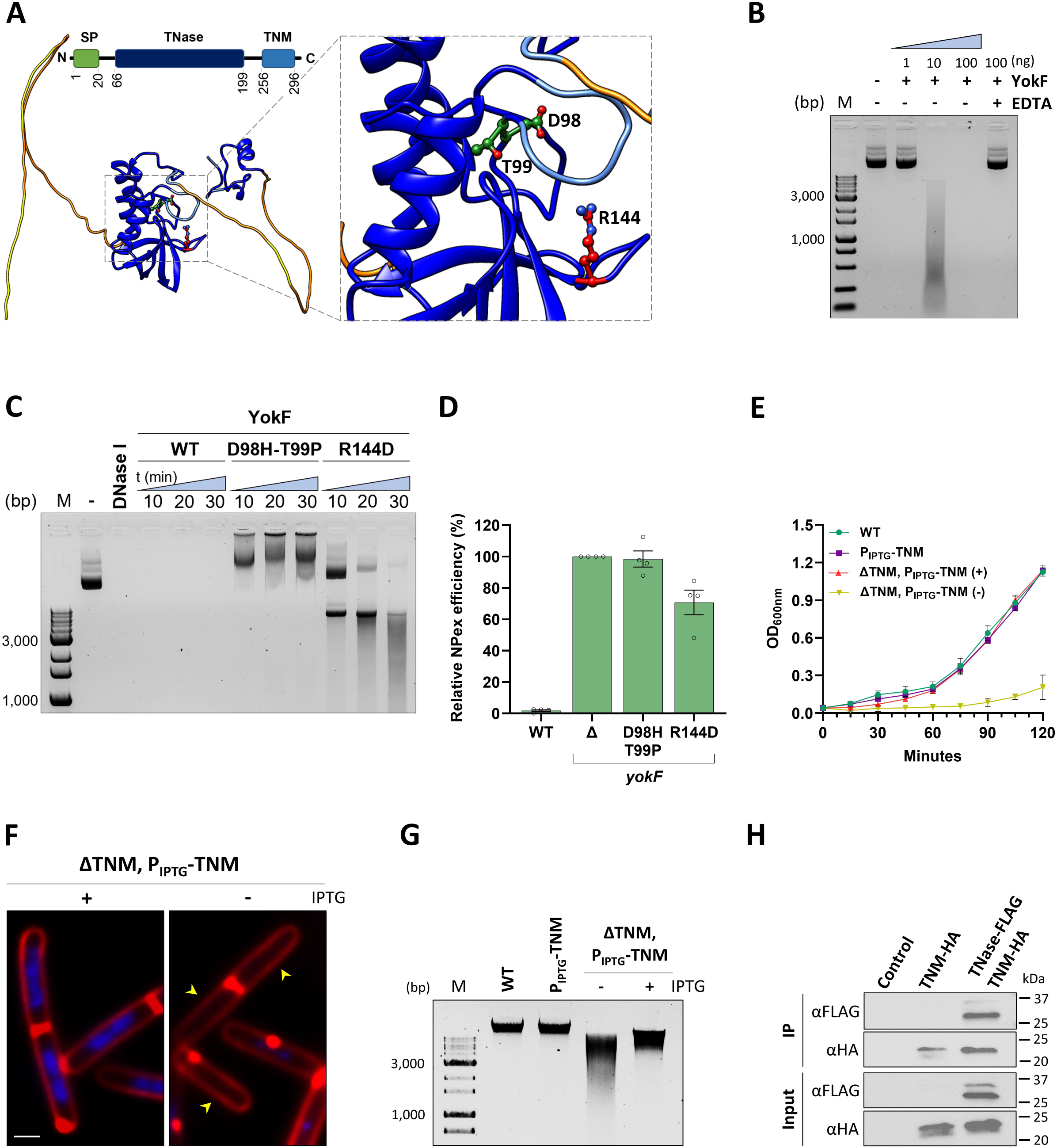
YokF displays nuclease activity crucial for NPex inhibition. **(A)** <u>YokF structure-function predictions</u>: Schematic representation and predicted AlphaFold structure of YokF illustrating its structural and functional domains. Schematics highlights the N-terminal signal peptide (SP: 1-20aa), thermonuclease domain (TNase: 66-199aa), and C-terminal TNM domain (TNM: 256-296aa), based on analyses from NCBI BLAST, SignalP 6.0, and EMBL-EBI InterPro. AlphaFold structure depicts the central TNase and the TNM domains with high pLDDT (predicted local difference test) value shown in dark blue ribbon, indicating a high level of confidence in the structural prediction. The inset box emphasizes the predicted active and binding sites clustered within the TNase domain, with the position of residues D98, T99, and R144, mutated in this study, highlighted. **(B)** <u>YokF displays nuclease activity *in vitro*</u>: Purified YokF at increasing concentrations was incubated with p6.6 kb (pHB201) at 37°C for 30 minutes, with or without EDTA. Shown is an agarose gel electrophoresis analysis of p6.6 kb DNA reaction products. M: Molecular weight marker [bp]. **(C)** <u>YokF functional analysis</u>: 100 ng of purified YokF (WT), YokF_D98H-T99P,_ or YokF_R144D_ were incubated with p6.6 kb (pHB201) at 37°C for 10, 20, and 30 minutes. DNase I reaction was carried out for 30 minutes in parallel. Shown is an agarose gel electrophoresis analysis of p6.6 kb DNA reaction products at the indicated time points. (-) no YokF. M: Molecular weight marker [bp]. **(D)** <u>YokF mutants are deficient in blocking NPex</u>: BSB1-derived donor strains: WT (BSB1) (GV345), Δ*yokF* (GV417: Δ*yokF*::*tet*), *yokF*_D98H-T99P_ (GV604: *yokF*_D98H-T99P_-*tet*), and *yokF*_R144D_ (GV580: *yokF*_R144D_-*tet*), carrying p6.6 kb (pHB201/*cat*, *erm*), were mixed in a 1:1 ratio with equivalent kanamycin resistance recipients: WT (GV372: *amyE*::P_IPTG_-*gfp*-*kan*), Δ*yokF* (GV416: Δ*yokF*::*tet*, *amyE*::P_IPTG_-*gfp*-*kan*), *yokF*_D98H-T99P_ (GV603: *yokF*_D98H-T99P_-*tet, amyE*::P_IPTG_*-gfp-kan*), and *yokF*_R144D_ (GV579: *yokF*_R144D_-*tet, amyE*::P_IPTG_*-gfp-kan*). The mixtures were incubated for 4 hours without selection, and equal volumes of cells were then spread over plates containing chloramphenicol, erythromycin, and kanamycin to select for trans-recipients. NPex efficiency was determined as the ratio of trans-recipients (CFU)/total recipients (CFU). Shown is NPex efficiency (%) relative to Δ*yokF* pair. Data is presented as mean values and SEM, based on 4 independent experiments. **(E)** <u>YokF features a self-inhibiting TNM domain</u>: Growth curves of BSB1-derived strains: YokF (WT), P_IPTG_-TNM (GV573: *amyE*::P_IPTG_*-yokF*_216-296aa_-*spec*), and ΔTNM, P_IPTG_-TNM (GV619: *yokF*_1-215aa_-*tet*, *amyE::*P_IPTG_-*yokF*_216-296aa_-*spec*). Cell growth, with (+) or without (-) IPTG, was followed by measuring OD_600nm_ at 15 minute intervals as indicated. Data is presented as mean values and SEM, based on 3 independent experiments. **(F)** <u>Deletion of the TNM domain results in chromosome degradation</u>: BSB1 cells expressing the YokF TNM domain under IPTG inducible promoter (ΔTNM, P_IPTG_-TNM) (GV619: *yokF*_1-215aa_-*tet*, *amyE::*P_IPTG_-*yokF*_216-296aa_-*spec*) were grown for 2 hours with (+) or without (-) IPTG. Shown are overlay images of cells stained with DAPI (blue) and FM4-64 (red). Arrows highlight cells lacking a signal from DAPI staining. Scale bar 0.5 µm. **(G)** <u>TNM domain inhibits TNase activity</u>: BSB1-derived strains: YokF (WT), P_IPTG_-TNM (GV573: *amyE*::P_IPTG_*-yokF*_216-296aa_-*spec*), and ΔTNM, P_IPTG_-TNM (GV619: *yokF*_1-215aa_-*tet*, *amyE::*P_IPTG_-*yokF*_216-296aa_-*spec*) were grown on LB plates with (+) or without (-) IPTG, and genomic DNA was extracted. Shown are equal amounts of genomic DNA samples subjected to agarose gel electrophoretic analysis. M: Molecular weight marker [bp]. **(H)** <u>TNase and TNM domains interact *in vivo*</u>: Lysates of BSB1-derived strains: WT (Control), TNM-HA, (GV627: *amyE*::P_IPTG_*-yokF_216-296aa_-HA_2_*_x_-*spec*), and TNase-FLAG, TNM-HA (GV638: *yokF*_1-215aa_*-FLAG_2_*_x-_*tet*, *amyE*::P_IPTG_-*yokF*_216-296aa_*-HA_2_*_x_-*spec*) were subjected to co-immunoprecipitation using anti-HA antibodies. Shown is a western blot analysis of the input and immunoprecipitated (IP) proteins using anti-HA or anti-FLAG antibodies, as indicated.

To test the prediction that YokF harbors nuclease activity, the purified protein was incubated with DNA. Indeed, DNA degradation was monitored in a YokF concentration-dependent manner (Fig. 2B). Structural docking over the TNase domain homologs located potential residues crucial for DNA binding and nuclease catalytic site (Fig. 2A; Fig. S3A)^52^. Accordingly, structure-directed mutations in YokF were generated: D98H-T99P, expected to abolish TNase catalytic activity, and R144D, predicted to affect YokF DNA binding (Fig. 2A; Fig. S3A). As anticipated, YokF_D98H-T99P_ lost its nuclease activity, causing a shift in DNA mobility due to the retained DNA binding (Fig. 2C). YokF_R144D_ exhibited partial deficiency in DNA degradation, likely due to reduced DNA binding (Fig. 2C). Subsequently, these mutations were introduced into the native YokF of BSB1 to assess their impact on NPex. YokF_D98H-T99P_ completely lost its NPex inhibition activity, while YokF_R144D_ showed a partial phenotype (Fig. 2D; Fig. S3A-S3B; Table S1). Collectively, we conclude that DNA binding and nuclease activities of the TNase domain are essential for NPex blockage by YokF.

Since YokF could be readily overexpressed without causing toxicity, we reasoned that its TNase activity is restrained *in vivo*, and unleashed only when required. Based on the predicted YokF structure, we hypothesized that the TNM domain might act to modulate the TNase activity. Consistently, the expression of YokF lacking the TNM domain (YokF_ΔTNM_) was toxic, though toxicity was relieved by co-expressing the TNM domain (Fig. 2E; Fig. S3C-3D). Ceasing TNM expression was associated with massive chromosome degradation, as indicated by DAPI staining and DNA extraction (Fig. 2F-2G; Fig. S3E). Correspondingly, when both domains were differentially tagged and co-expressed *in vivo*, immunoprecipitation analysis demonstrated their direct interaction (Fig. 2H). Taken together, YokF nuclease activity is modulated *in vivo* through association with the TNM domain.

### Membrane localization of YokF is required for NPex

We next investigated YokF subcellular localization by immunofluorescence microscopy using anti-YokF antibodies. The analysis revealed the protein to be localized as foci decorating the cell membrane, consistent with its predicted SP (Fig. 3A). YokF-GFP fusion did not yield a detectable signal (Fig. 3B), suggesting that the C-terminal GFP could be secreted or cell surface located ^53^. In agreement, removing the YokF SP domain turned the GFP fusion visible, shifting the protein localization to the cytoplasm (Fig. 3B). Importantly, YokF_ΔSP_ lost its ability to inhibit NPex (Fig. 3C), indicating the necessity of membrane localization for its activity. Biochemical fractionation analysis using anti-YokF antibodies unveiled the full-length protein (∼37 kDa) alongside a faint shorter ∼25 kDa fragment in both cytoplasmic and membrane fractions, with the membrane localization being dependent on the SP (Fig. 3D). Intriguingly, a secreted SP-dependent fragment of YokF (∼18 kDa) was detected (Fig. 3D). Tagging YokF C-terminus with HA enabled the detection of the secreted fragment, specifying that the C-terminus TNM domain was released into the medium (Fig. 3D). We surmise that the TNM domain of membrane-localized YokF is removed to locally relieve its TNase activity, a crucial step required for NPex prevention.

**Fig. 3.**
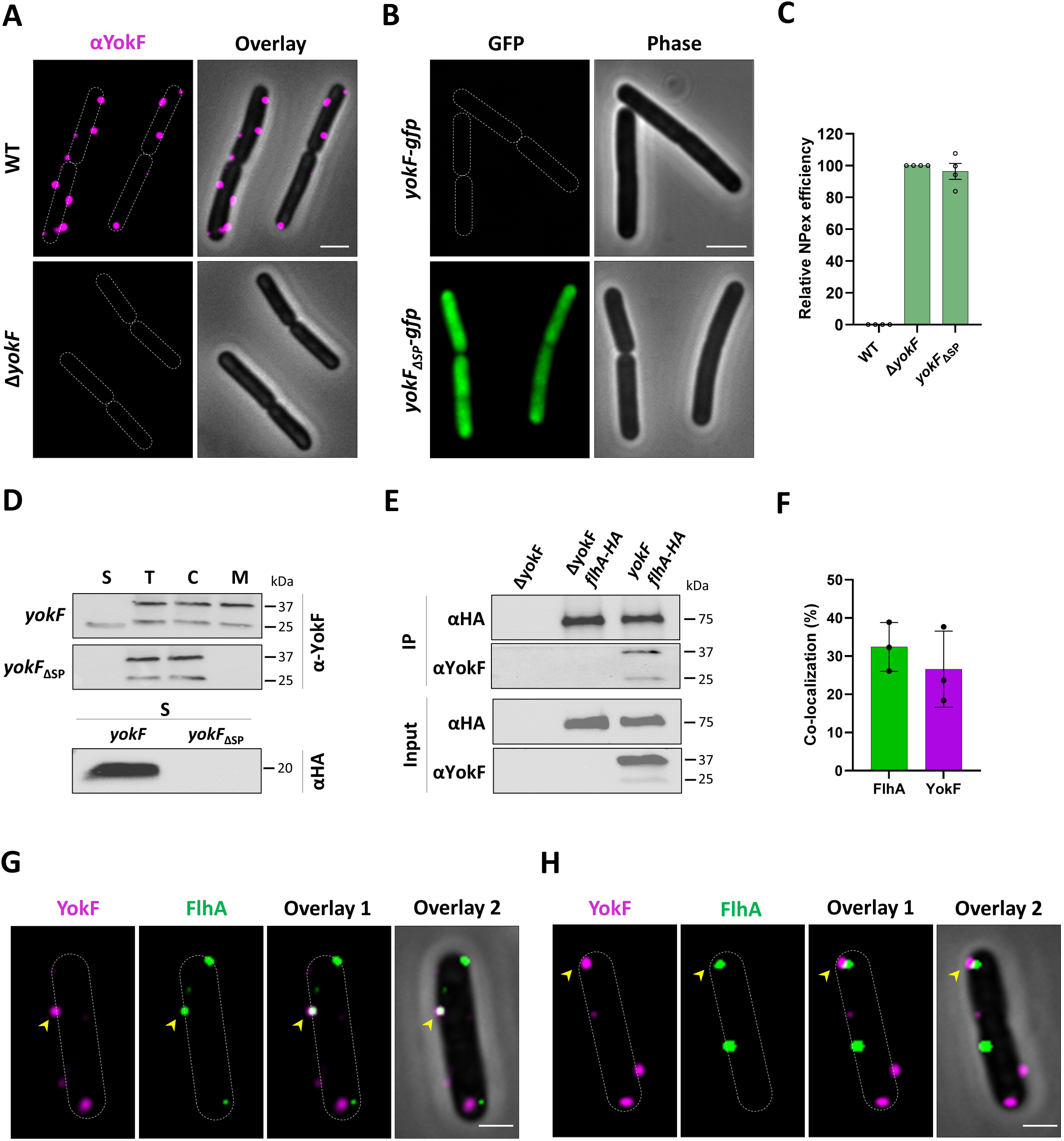
YokF is a membrane-localized protein that interacts with FlhA. **(A)** <u>YokF localizes to the membrane</u>: BSB1 (WT) or isogenic Δ*yokF* (GV411: Δ*yokF*::*tet*) strains were treated with anti-YokF primary antibodies followed by Alexa 647-conjugated secondary antibodies, and subjected to immunofluorescence microscopy. Shown are immunofluorescence images, with highlighted cell borders (αYokF, left), and their overlay with corresponding phase contrast (gray) images (right). Scale bar 1 µm. **(B)** <u>YokF_ΔSP_ shows cytoplasmic localization</u>: BSB1-derived strains harboring YokF-GFP (GV475: *yokF*-*gfp*-*kan*) or YokF_ΔSP_-GFP (GV483: *yokF*_ΔSP_-*gfp*-*tet*) were visualized by fluorescence microscopy. Shown are fluorescence images from GFP (left), with highlighted cell borders (upper panel), and corresponding phase contrast images (right). Scale bar 1 µm. **(C)** <u>SP domain is required for NPex</u>: BSB1-derived donor strains: WT (BSB1) (GV345), Δ*yokF* (GV417: Δ*yokF*::*tet*), and *yokF*_ΔSP_ (GV486: *yokF*_ΔSP_-*tet*), carrying p6.6 kb (pHB201/*cat*, *erm*), were mixed in a 1:1 ratio with equivalent kanamycin resistance recipients: WT (GV372: *amyE*::P_IPTG_-*gfp*-*kan*), Δ*yokF* (GV416: Δ*yokF*::*tet*, *amyE*::P_IPTG_-*gfp*-*kan*), and *yokF*_ΔSP_ (GV485: *yokF*_ΔSP_-*tet, amyE*::P_IPTG_*-gfp-kan*). The mixtures were incubated for 4 hours without selection, and equal volumes of cells were then spread over plates containing chloramphenicol, erythromycin, and kanamycin to select for trans-recipients. NPex efficiency was determined as the ratio of trans-recipients (CFU)/total recipients (CFU). Shown is NPex efficiency (%) relative to Δ*yokF* pair. Data is presented as mean values and SEM, based on 4 independent experiments. **(D)** <u>TNM domain is secreted</u>: Upper panels: BSB1-derived strains expressing native *yokF* (WT) or *yokF*_ΔSP_ (GV484: *yokF*_ΔSP_-*tet*) were grown to mid-logarithmic phase and proteins were extracted from total lysate (T), cytoplasmic (C), membrane (M), and supernatant (S) fractions. Shown is a western blot analysis using anti-YokF antibodies. Lower panel: Supernatant of BSB1-derived strains *yokF* (GV455: *yokF-HA*_x2_-*tet*), and *yokF*_ΔSP_ (GV487: *yokF*_ΔSP_*-HA*_x2_-*tet*) tagged with HA were subjected to a western blot analysis using anti-HA antibodies. **(E)** <u>YokF interacts with FlhA</u>: Lysates of BSB1-derived strains: Δ*yokF* (GV411: Δ*yokF*::*tet*), Δ*yokF, flhA-HA* (GV271: Δ*yokF*::*tet, flhA*-*HA*_x2_*-kan*, P*_fla/che_*-*flhF*), and *yokF, flhA-HA* (GV272: *flhA*-*HA*_x2_*-kan*, P*_fla/che_*-*flhF*) were subjected to co-immunoprecipitation using anti-HA antibodies. Shown is a western blot analysis of the input and immunoprecipitated (IP) proteins using anti-HA or anti-YokF antibodies, as indicated. **(F)** <u>YokF and FlhA colocalize</u>: BSB1-derived strain expressing FlhA-SG (GV617: *flhA-lin_4_*_x_*-sg_d_-kan*-P*_fla/che_*-*flhF*) was treated with anti-YokF primary antibodies followed by Alexa 647-conjugated secondary antibodies and subjected to fluorescence microscopy. Shown is quantitative representation (%) of FlhA foci that colocalized with YokF (green), and YokF foci that colocalized with FlhA (magenta) (n > 300 cells). SD was calculated for different fields. A representative experiment out of 3 independent repeats. **(G-H)** <u>Representative fluorescence images highlighting YokF and FlhA colocalization</u>: Shown are YokF (magenta), FlhA-SG (green), their overlay (Overlay 1), and their overlay with phase contrast (gray) (Overlay 2). Arrows highlight YokF and FlhA colocalization sites, exemplifying full (G), and partial colocalization (H). Scale bars 0.5 µm.

To uncover YokF interacting partners, we conducted immunoprecipitation coupled with mass spectrometry (MS), utilizing YokF as bait (Table S2). Among the strong interactors identified was FlhA, a pivotal CORE component, mediating NPex^26^. Subsequently, immunoprecipitation followed by western blot analysis substantiated the existence of such interaction (Fig. 3E). Co-visualization of YokF and FlhA by fluorescence microscopy showed frequent colocalization of both protein foci (Fig. 3F-3H; Fig. S4A-S4B). As YokF expression does not abolish nanotube formation (Fig. S2A), we infer that membrane-localized YokF interacts with FlhA, functioning as a gatekeeper to inhibit DNA delivery via nanotubes.

### YokF represents a widespread family of NPex inhibitors

To expand the role of YokF, we surveyed for its potential conservation among bacteria. Notably, *B. subtilis* carries YncB, showing homology to YokF SP and TNase domains but lacking TNM (Fig. S5A). Deletion of *yncB* had no impact on NPex levels (Fig. S5B-S5D), insinuating that YokF functional homologs should comprise all three domains. Through a conservation analysis, we identified YokF homologs in a wide range of Gram-positive bacteria (Table S3) and compiled a dataset of 44 homologs representing phylogenetic diversity. The examination revealed YokF homologs in various *Bacillaceae, Staphylococcaceae,* and other Gram-positive families (Fig. 4A). Further bioinformatic analysis revealed that apart from *B. subtilis* and its closest species *B. amyloliquefaciens*, all other investigated YokF homologs were not located in obvious prophage regions, and appeared as part of the host chromosome (Fig. 4A; Table S3), signifying that NPex inhibition may be a genuine host-associated trait.

**Fig. 4.**
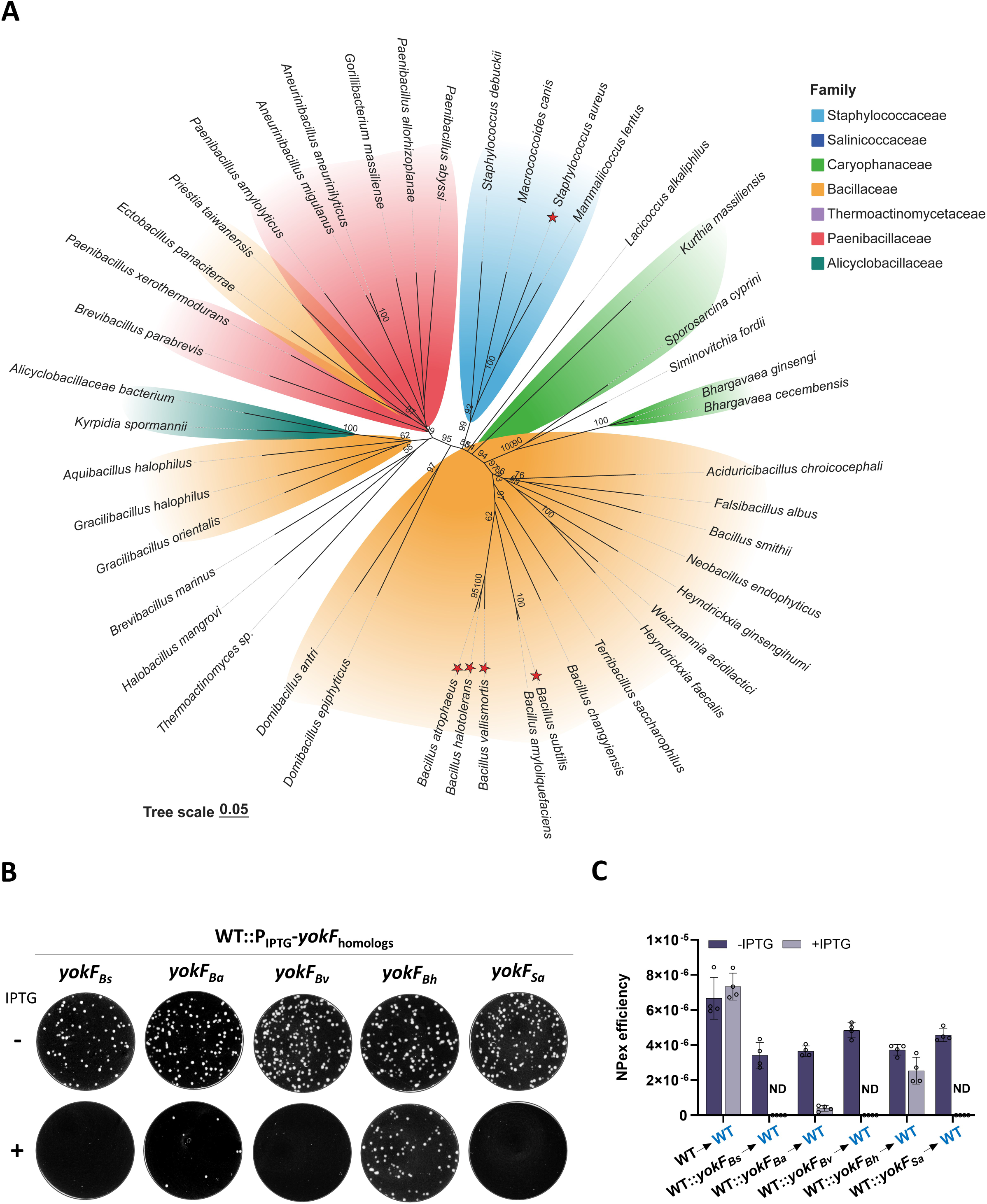
A widespread family of YokF homologs. **(A)** <u>YokF is highly conserved among Gram-positive bacteria</u>: Homology sequences of YokF were collected from the NCBI protein databases (Table S3). Sequences were subsequently aligned via the ClustalW methodology^63^. The phylogenetic trees of YokF homologs were constructed using the neighbor-joining algorithm of MEGA 7.0 with 1000 bootstrap values, and displayed with an online platform TVBOT v2.6. Bootstrap values greater than 50 are indicated at the nodes. Major families are highlighted in different colors as specified. Scale bar represents 0.05 substitutions per site. Red stars indicate homologs tested for their functionality. **(B)** <u>YokF homologs inhibit NPex</u>: PY79-derived donor strains, harboring YokF homologs from *B. subtilis* (BSB1) (*Bs*) (GV480: *amyE*::P_IPTG_*-yokF_Bs_*-*spec*), *B. atrophaeus* (*Ba*) (GV653: *amyE*::P_IPTG_*-yokF_Ba_*-*spec*), *B. vallismortis* (*Bv*) (GV655: *amyE*::P_IPTG_*-yokF_Bv_*-*spec*), *B. halotolerans* (*Bh*) (GV657: *amyE*::P_IPTG_*-yokF_Bh_*-*spec*), and *S. aureus* (*Sa*) (GV659: *amyE*::P_IPTG_*-yokF_Sa_*-*spec*), all carrying the p6.6 kb (pHB201/*cat*, *erm*), were grown with (+) or without (-) IPTG, and mixed in a 1:1 ratio with kanamycin resistance WT recipient (GV236: *sacA*::*kan*). The mixtures were incubated for 4 hours without selection, in the presence (+) or absence (-) of IPTG. Equal volumes of cells were then spread over plates containing chloramphenicol, erythromycin, and kanamycin to select for trans-recipients. Shown are representative images of NPex selective plates. **(C)** Quantitative analysis of NPex efficiencies between the donor and recipient pairs shown in (B). NPex efficiency=Trans-recipients CFU/Total recipients CFU. Data are presented as mean values and SEM, based on 4 independent experiments. ND – Not Detected.

To decipher whether the anti-NPex YokF function is shared across species, we introduced into *B. subtilis* (PY79) *yokF* homologs from closely related *Bacilli*, *B. vallismortis* (YokF-*_Bv_*), *B. halotolerans* (YokF*-_Bh_*), and *B. atrophaeus* (YokF-*_Ba_*) (Fig. S6A), and assessed their NPex inhibition efficiency. Remarkably, YokF-*_Ba_* and YokF-*_Bv_* fully blocked NPex upon induction, whereas YokF-*_Bh_* showed partial activity (Fig. 4B-4C), with none of these homologs exhibiting a significant effect on *B. subtilis* growth (Fig. S6B). Motivated by these findings, we next tested, in a similar manner, a YokF homolog derived from the evolutionary distant *Staphylococcus aureus* (YokF-*_Sa_*) pathogen. Despite lower identity (39%), YokF-*_Sa_* effectively blocked *B. subtilis* NPex (Fig. 4B-4C; Fig. S6A-S6B). Thus, we conclude that YokF anti-NPex activity is prevalent among Gram-positive bacteria and is likely to modulate the spread of non-conjugative plasmids in natural settings.

## Discussion

Here we report that bacterial intercellular nanotubes play a significant role in plasmid dissemination, functioning as vehicles for plasmid donation and, to a reduced extent, plasmid acquisition. The widespread presence of nanotubes across diverse eubacteria and archaebacteria [e.g.,^23,28–34,36^] emphasizes the potential contribution of this pathway to horizontal gene transfer. Through the identification of a *B. subtilis* strain impaired in NPex, we discovered YokF, a membrane-located TNase, as a prophage-derived factor that actively inhibits plasmid spread. Unlike known plasmid antagonistic systems, such as restriction enzymes, that operate from the recipient side^54,55^, YokF primarily impedes plasmid donation to neighboring bacteria. Based on our findings, we propose that membrane-localized YokF becomes activated through the removal of its autoinhibitory TNM domain. Its intimate interaction with the nanotube platform component FlhA locates its degradation activity to effectively block the passage of DNA into the channel, serving as a conduit gatekeeper that controls plasmid flow (Fig. 5). The existence of YokF in the genome of many Gram-positive bacteria, along with their shared functionality demonstrated here, reflects on their significant role in modulating plasmid dispersal in nature and hints that additional such factors are likely to be abundant. The utilization of YokF-like factors by the host may allow restricting the spread of plasmids, harboring beneficial genes, to competing non-kin bacteria.

**Fig. 5.**
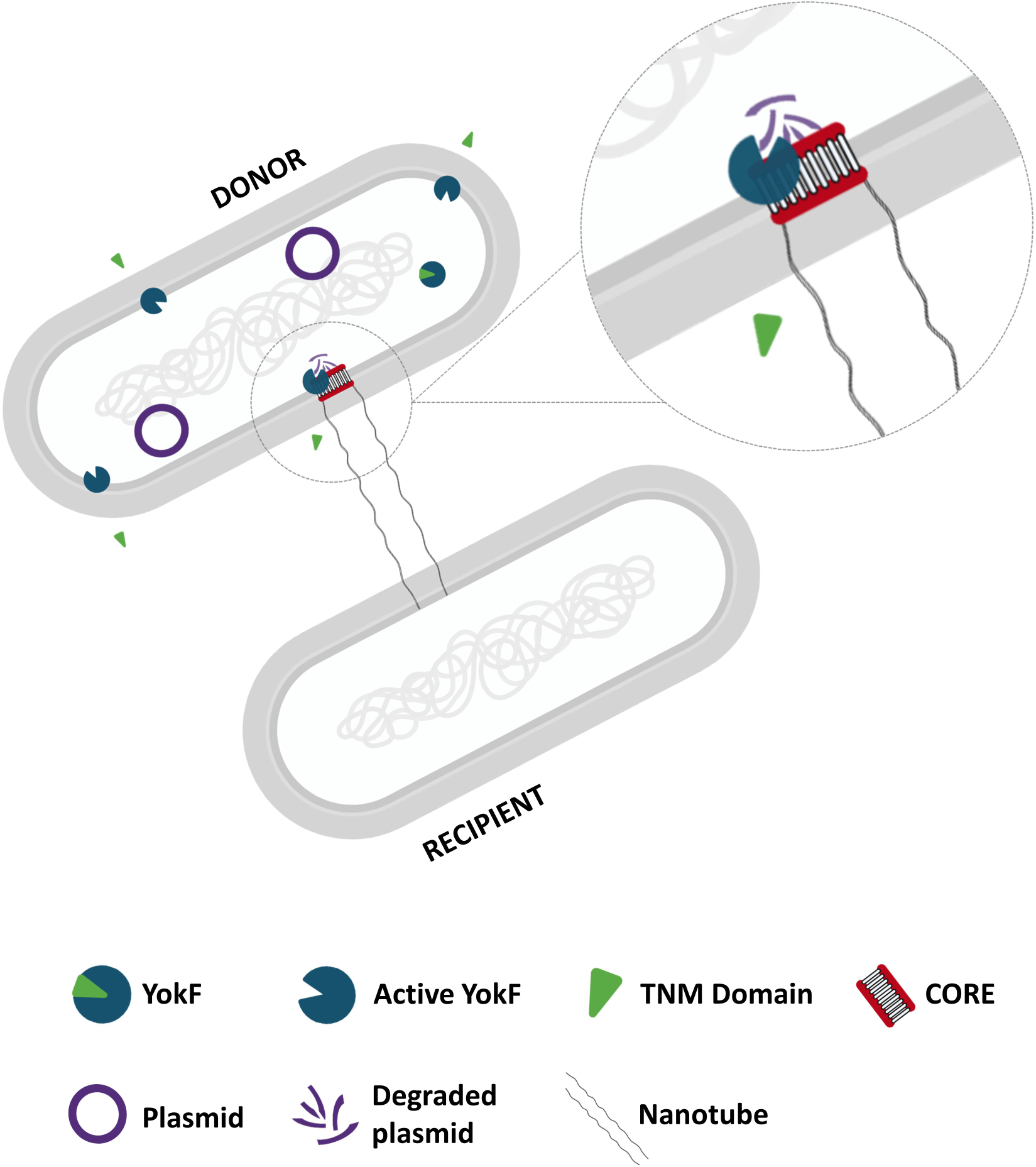
A model for YokF anti-NPex activity. Schematic illustrating the mechanism by which YokF inhibits NPex. Following membrane localization, YokF undergoes a proteolytic cleavage, leading to TNM secretion and TNase activation. Activated YokF interacts with the CORE component FlhA, degrading plasmid DNA in the nanotube vicinity, thereby impeding NPex. An enlarged view shows YokF action site.

In *B. subtilis* YokF is encoded by the SPβ prophage^49,56^, while most of YokF homologs were found to be integral constituents of the host genome, suggesting that SPβ may have acquired *yokF* through transduction. The ability of mobile genetic elements to exclude incoming DNA reflects a broader phenomenon. Several plasmids, for instance, encode factors that restrict the host competence machinery. A non-domesticated *B. subtilis* isolate, encodes a plasmid-borne small protein ComI, which inhibits the competence machinery^57^, whereas diverse conjugative elements in the *Legionella* genus carry small RNAs capable of silencing natural transformation^58^. Furthermore, plasmids of pathogenic *Vibrio cholerae* strains possess systems to degrade and eliminate small multi-copy plasmids^59–62^. These findings reflect on the complexity of plasmidome dynamics, with YokF being a key factor in NPex modulation in Gram-positive bacteria. The realization of NPex as a potential driver of horizontal gene transfer, beyond the classical conjugation, transformation, and transduction, underscores the existence of additional overlooked non-canonical DNA transfer mechanisms that operate in nature.

## Acknowledgments

We thank the members of the Ben-Yehuda and Rosenshine laboratories for their insightful discussions. We are also grateful to David Rudner (Harvard University), Oscar P. Kuipers (University of Groningen), and Libor Krasny (Academy of Sciences of the Czech Republic) for providing bacterial strains and plasmids. We thank Manoj Kumar (Hebrew University), and Abed Nasereddin (Core Research Facilities, Hebrew University) for technical support. VG was supported by the Planning and Budgeting Committee (PBC) of Israel postdoctoral fellowship, and SB by the Golda Meir postdoctoral fellowship.

## Author contributions

Conceptualization: VG, IR, and SB-Y

Methodology: VG, SB, MBS, MZ, BS, MKG, YN, TK, IR, and SB-Y

Investigation: VG, SB, MBS, MZ, BS, IR, and SB-Y

Visualization: VG, SB, MBS, MZ, BS, MKG, IR, and SB-Y

Funding acquisition: SB-Y, and IR

Project administration: SB-Y, MR, and IR

Supervision: SB-Y, and IR

Writing - original draft: VG, SB-Y, and IR

## Data and materials availability

All data are available in the main text or the supplementary materials.

## Supplementary Materials

### Methods

#### Bacterial strains and plasmids

Bacterial strains, plasmids, and primers utilized in this study are listed in Tables S4 and S5, respectively. Antibiotic resistance cassettes flanked by lox66/lox71 sites were amplified from pWX465 (cat), pWX466 (spec), pWX467 (erm), pWX469 (tet), and pWX470 (kan) using the primers 2430 and 2431. Plasmid constructions were carried out in *Escherichia coli* DH5α using standard molecular biology techniques^64^.

#### General growth conditions

All general procedures were conducted as previously described ^65^. Briefly, *B. subtilis* cultures were inoculated at OD_600nm_ 0.05 from overnight cultures and grown at 37°C in LB medium (Difco). For gene induction under the P_IPTG_ promoter, 0.5-1 mM Isopropyl-β-D-thiogalactopyranoside (IPTG) was used. Selection of *B. subtilis* strains was performed with the following antibiotic concentrations: kanamycin (10 µg/ml, US Biological), chloramphenicol (5 µg/ml, Sigma-Aldrich), lincomycin (25 µg/ml, Sigma-Aldrich), erythromycin (1 µg/ml, Sigma-Aldrich), tetracycline (10 µg/ml, Sigma-Aldrich), spectinomycin (100 µg/ml, Sigma-Aldrich). For *E. coli* strains, the selection was conducted using ampicillin at 100 μg/mL, and kanamycin at 40 μg/mL.

#### Site-directed mutagenesis

DNA fragments were amplified using YokF-SDM primers (7017-7022 and 7282-7285) (Table S5). The fragments were assembled using a Gibson assembly mix (NEB), and then transformed into the indicated *B. subtilis* strains. Point mutations were verified by DNA sequencing.

#### NPex efficiency determination

An NPex efficiency assay was conducted to detect plasmid exchange. Donor cells carrying a plasmid, and recipient strains lacking a plasmid were grown separately to OD_600nm_ 0.8, after which they were mixed in a 1:1 ratio. The mixtures were incubated in LB medium, without antibiotic selection, at 37°C with gentle shaking (35 rpm) for 4 hours. 0.5 mM IPTG was added when indicated. After incubation, equal numbers of cells were plated onto double- or triple-selective LB agar plates containing the relevant antibiotics to select recipient cells that had acquired the plasmid (termed, trans-recipients). Concurrently, cells were serially diluted and plated onto either kanamycin or lincomycin plates to quantify the total CFU of recipient bacteria, and onto LB plates, without antibiotics, as a control. NPex efficiency was calculated as the ratio of the number of trans-recipients CFU/ total recipient CFU.

#### Protein exchange assay

Exchange of antibiotic-modifying enzymes (i.e., chloramphenicol acetyltransferase and kanamycin resistance protein), was assayed as previously described ^4^. Briefly, donor and recipient strains (Table S4) were grown to OD_600nm_ 0.8, mixed in a 1:1 ratio, and incubated in LB medium, without antibiotic selection, at 37°C with gentle shaking for 4 hours. After incubation, equal numbers of cells were spotted (10 µl) on double-selective LB agar plates containing chloramphenicol (6 µg/ml) and kanamycin (5 µg/ml). As a control, cells were also spotted on LB agar without antibiotics. Plates were incubated overnight at 37°C and imaged over time.

#### Motility assay

The motility assay was conducted as previously described^66^ with some modifications. Briefly, cells were grown to the mid-logarithmic phase in LB and then concentrated 10-fold to an OD_600nm_ 5.0. A 5 μl aliquot of the cell suspension was spotted onto freshly prepared LB plates containing 0.3% agar. The plates were incubated for 18 hours at 37°C and imaged over time using a photo scanner (EPSON-V800).

#### Plasmid spread assay

Donor strains, PY79 (WT) or PY79 expressing YokF, carrying p6.6 kb (pHB201/*cat*, *erm*), and a PY79-derived recipient strain (Δ*comK*, Δ*CORE*), were grown separately to OD_600nm_ 0.8, mixed in a 1:1 ratio, and incubated in LB medium, without antibiotic selection, at 37°C with gentle shaking for 4 hours. 0.5 mM IPTG was added when indicated. Next, equal numbers of cells from the co-culture were spotted on triple-selective LB agar plates containing chloramphenicol, kanamycin, and lincomycin to detect plasmid exchange. Concurrently, cells were serially diluted and plated onto kanamycin-containing plates to quantify the total CFU of recipient bacteria (designated day 0). The following day, the recipient population was collected from the kanamycin plates (10^0^ dilution), suspended in fresh LB medium, and grown to the mid-logarithmic phase. These cells were then mixed with fresh mid-log donor cells in a 1:1 ratio, and incubated for 4 hours at 37°C with gentle shaking under mild selective pressure (5µg/ml Lincomycin), without causing cell death. This process was repeated for 8 consecutive cycles. The plasmid dissemination rate was calculated as the ratio of the number of recipients that acquired the plasmid (CFU)/ total recipient (CFU) on each day.

#### Plasmid stability assay

Bacteria harboring p6.6 kb (pHB201/*cm*, *erm*), grown overnight with antibiotic selection (0^th^ generation), were diluted to OD_600nm_ 0.05 in fresh LB medium, without antibiotics, and incubated at 37°C with gentle shaking for approximately 10 generations, as estimated by optical density. Every 10 generations, cultures were back-diluted to OD_600nm_ 0.05 and continued to grow up to ∼30 generations. Cultures withdrawn at each 10^th^ generation, including generation 0, were serially diluted and plated on LB with and without erythromycin. Plates were incubated overnight at 37°C. Plasmid stability was determined by calculating the ratio of the number of erythromycin-resistant colonies (indicative of plasmid retention) (CFU)/total population (CFU).

#### Transformation efficiency assay

Fresh colonies were suspended in 1 ml of modified competence medium (1xMC) [80 mM K_2_HPO_4_, 30 mM KH_2_PO_4_, 2% Glucose, 30 mM Trisodium citrate, 22 µg/ml Ferric ammonium citrate, 0.1% Casein Hydrolysate (CAA), 0.2% potassium glutamate], and grown under shaking conditions at 37°C for 3 hours. Cultures were then adjusted to OD_600nm_ 0.8 with 1xMC medium and supplemented with 100 ng of either plasmid or genomic DNA (gDNA), followed by an additional 3 hour incubation at 37°C. To select for transformants, 100 µL of the cell suspension was plated onto LB agar plates containing the appropriate antibiotics. In parallel, serial dilutions of cell suspension were plated on LB without antibiotics to enumerate the total population. Transformation efficiency was calculated as the ratio of the number of transformants (CFU)/total population (CFU).

#### Assessing the contribution of transduction and membrane vesicles to NPex

Donor cells carrying a plasmid, and recipient cells lacking a plasmid were grown separately to an OD_600nm_ 0.8. Donors and recipient strains were mixed in a 1:1 ratio or cultured separately in fresh LB medium, without antibiotic selection, and incubated at 37°C with gentle shaking (35 rpm) for 3 hours. Following incubation, cells were gently centrifuged, and the donor culture supernatant was collected, filtered through a 0.22 µm membrane, and used to replace the medium of the recipient strain. The recipients were subsequently incubated with the donor supernatant at 37°C with gentle shaking (35 rpm) for 1 hour. After incubation, equal numbers of cells were plated onto triple-selective LB agar plates containing the relevant antibiotics to select recipient cells that had acquired the plasmid (termed, trans-recipients). Concurrently, cultures were serially diluted and plated onto kanamycin plates to quantify the total CFU of recipient bacteria, and onto LB plates, without antibiotics, as a control. Plasmid transfer efficiency was calculated as the ratio of trans-recipients CFU to total recipient CFU.

#### Plaque assay

Overnight cultures of bacterial strains were refreshed to the mid-logarithmic phase and infected with SPO1 or SPP1 phages and incubated for 12 minutes at 37°C to facilitate phage attachment. Following infection, cultures were plated onto 1.5% MB (LB supplemented with 5mM MgCl_2_ and 0.5mM MnCl_2_) agar plates and further incubated at 37°C for 18-20 hours.

#### Nanotube visualization

Nanotube visualization was carried out as previously described^44^. Accordingly, cells grown to the mid-logarithmic phase were spotted onto EM grids (mesh copper grids, EMS) placed over LB agar plates and incubated for 4 hours at 37°C. Cells were washed 3 times with 1xPBS, fixed with 2% paraformaldehyde and 0.01% glutaraldehyde in sodium cacodylate buffer (0.1 M, pH 7.2) for 10 minutes at 25°C. Cells were left overnight for fixation in 2% glutaraldehyde in sodium cacodylate buffer (0.1 M, pH 7.2) at 4°C. EM grids underwent a series of washes for cell dehydration in increasing ethanol concentrations (25, 50, 75, and 96%), and were kept in a vacuum until visualization. Samples were coated with iridium (Safematic), and observed using an In-Lens detector operated at Secondary Electron (SE) mode by Gemini 560 HR SEM (Ziess).

#### Fluorescence microscopy

For visualization of YokF-GFP, exponentially growing cells were harvested at OD_600nm_ 0.5, washed with 1xPBS, and observed by fluorescence microscopy. For FM4-64 and DAPI staining, cells were harvested at an OD_600nm_ 0.5, washed with 1XPBS, and resuspended in 50 μL of 1XPBS containing FM4-64 (5 µg/mL, Thermo Fisher) or DAPI (5 µg/mL, Sigma), and visualized using ECLIPSE Ti2 microscope (Nikon, Japan), equipped with Prime BSI camera (Photometrics, Roper Scientific, USA). System control and image analysis were performed using NIS-Elements AR Analysis (version 5.30.07, Nikon, Japan).

#### Immunofluorescence microscopy

Immunofluorescence microscopy was carried out as previously described^67^ with some modifications. Overnight cultures were diluted in fresh LB medium to OD_600nm_ 0.1 and incubated for 2 hours at 37°C with gentle shaking. To fix the cells, 1 ml of the culture was mixed with 10 ml of ice-cold methanol (80%) and incubated at room temperature for 1 hour. Subsequently, 200 µl of paraformaldehyde (0.3% final concentration) was added, and the mixture was incubated for 5 minutes at room temperature. Cells were then centrifuged at 3500 rpm, resuspended in 1 ml ice-cold water, and permeabilized with lysozyme (2 mg/ml) in GTE buffer [25 mM Tris, 50 mM glucose, 10 mM EDTA] for 30 seconds. Following permeabilization, the cells were washed twice with 1xPBS. For immunostaining, cells were incubated with anti-YokF primary antibodies (1:1000 in 2% BSA) for 30 minutes, washed thrice with 1xPBS, and then treated with Alexa Fluor 647 (Thermo Fisher Scientific) secondary antibodies (1:1000 in 2% BSA). After additional extensive washing with 1xPBS, cells were spotted onto Poly-L-Lysine-coated coverslips and visualized using fluorescence microscopy.

#### Protein purification and production of YokF polyclonal antibodies

*E. coli* BL21 (DE3, Invitrogen) cells harboring pGV1, pGV16, or pGV17, each expressing a different derivative of YokF tagged with His_x6_, were grown in LB medium supplemented with kanamycin (40 μg/mL), and protein synthesis was induced at OD_600nm_ 0.8 (0.5 mM IPTG, 10 hours, 30°C). Cell culture (1 L) was pelleted (8,000 rpm, 10 minutes, 4°C), resuspended (4 ml/gram wet weight) in lysis buffer [(50 mM NaH_2_PO_4_, 300 mM NaCl, 10 mM imidazole, pH 8.0), and PMSF protease inhibitor X1 (Sigma)], and lysed using an M-110S Microfluidizer (Microfluidics). The lysate was centrifuged (12,000 rpm, 45 minutes, 4°C), and the supernatant was filtered through a 0.2 µm syringe filter (Sartorius). The cleared extract was subjected to metal affinity chromatography (HisTrap HP column, Cytiva) using the ӒKTA ‘go’ (Cytiva) with a HisTrap HP column (Cytiva). The eluted samples were subjected to size exclusion chromatography (Superdex 200), and the pooled fractions were concentrated using centrifugal filters (Amicon Ultra-15). Purified proteins were stored at −80⁰C for further studies. Rabbit polyclonal antibodies against the purified YokF protein were generated by Eurogentec (Belgium) custom polyclonal antibody production service. In brief, rabbits were immunized with the purified YokF protein to induce an immune response. Following immunization, YokF-specific antibodies were purified through affinity chromatography. The initial validation of the purified antibodies was performed by Eurogentec, using a qualitative indirect enzyme-linked immunosorbent assay (ELISA).

#### Nuclease activity

Purified YokF (wild-type or mutant) was incubated with either plasmid DNA or gDNA (100 ng/µl) in a reaction buffer composed of 20 mM MES, 100 mM NaCl, 2 mM CaCl_2_, and 2 mM MgCl_2_, at pH 6.9. Bovine DNase I (1 µg/µL, Merk) (New England Biolabs) served as the positive control. The nucleic acid hydrolysis reaction was conducted at 37°C for 10-30 minutes, as indicated, and was then halted by EDTA addition to a final concentration of 5 mM. The integrity of the tested DNA was assessed by performing agarose gel electrophoresis (0.8-1% gel, 40 minutes at 100V), stained with ethidium bromide, and imaged using GelDoc Imaging System (Bio-Rad).

#### Protein extraction and western blot analysis

Overnight cultures were refreshed with fresh LB to OD_600nm_ 0.05 and incubated at 37°C with gentle shaking until reaching OD_600nm_ 1. The culture supernatant was collected, concentrated using Amicon filters (10 kDa cutoff), and supplemented with Halt™ protease-phosphatase inhibitor cocktail (1x, Thermo Scientific). Harvested cells were washed with 1XPBS and resuspended (2 ml/gram wet weight) in a lysis buffer containing 150 mM NaCl, 50 mM Tris-HCl (pH 8.0), 1.0% NP-40, lysozyme (1 mg/ml), and 1x protease inhibitor. Cell lysis was performed using FastPrep-24™ (MP) (level 4, 40 sec, ×4). The suspension was then centrifuged (15,000 rpm, 10 minutes, 4°C), and the supernatant was collected for total lysate. The collected fraction was further subjected to ultracentrifugation (55,000 rpm, 2 h, 4°C, Beckman LT100) to separate the cytoplasmic and membrane fractions. Proteins were subjected to SDS-PAGE using Mini-PROTEAN TGX Stain-Free Precast Gels (Bio-Rad), and electroblotted onto a nitrocellulose membrane with the Trans-Blot Turbo Transfer System (Bio-Rad). The membrane was blocked for 1 hour with 5% skim milk in TBST (50 mM Tris-Cl, pH 7.5, 150 mM NaCl, 1% Tween-20). Blots were then probed with either polyclonal rabbit anti-YokF, anti-HA, or monoclonal mouse anti-FLAG antibodies (1:5000, 0.05% Tween-20, 5% skim milk in TBS) followed by peroxidase-conjugated goat anti-rabbit or anti-mouse secondary antibody (Bio-Rad) (1:10,000, 0.05% Tween-20, 5% skim milk). The signal was detected using an EZ-ECL kit (Biological Industries, Beit Haemek, Israel). Gels and blots were imaged using the ChemiDoc MP imaging system (Bio-Rad). A stain-free image was used as total protein loading control.

#### Co-immunoprecipitation

Cultures grown to mid-logarithmic phase were collected by centrifugation (4,000 rpm, 10 minutes, 4°C), washed 3 times with 1X PBS buffer, pellets were resuspended in lysis buffer (1X PBS, 0.05% Tween 20, 10 mM Imidazole, 2X Protease inhibitor, 1 mM DTT), and lysed using a (MP) (level 4, 40 sec, ×4). Cells were cross-linked with paraformaldehyde (0.5% final concentration) for co-immunoprecipitation of YokF domains. The lysates were centrifuged (15,000 rpm, 10 minutes, 4°C), and total protein concentration was determined by Bradford dye-binding method with a protein assay kit (Bio-Rad). Clarified lysates with equal amounts of protein concentrations (1 ml) were incubated overnight with 30 µl of anti-HA magnetic bead resin (Sigma) (pre-washed with 1X TBST buffer) on an orbital shaker at 4°C. The beads were separated magnetically, washed with a washing buffer (1X TBS, 150 mM NaCl, 0.05% Tween 20, 10 mM Imidazole, 1 mM DTT), and processed for either Western blot or mass spectrometry (MS) analysis.

For Western blot analysis, 50 µl of 2X Laemmli sample buffer was added to the resin, followed by heating at 95°C for 10 minutes. After gentle vortexing and magnetic separation, the supernatant was transferred to a fresh tube for analysis. For MS analysis, the trypsin-digestion process was performed as described ^68^. The above-enriched resin was resuspended in 100 µl of denaturation buffer (6 M Urea, 2 M thiourea in 10 mM Tris, pH 8.0). Samples were reduced with DTT (final concentration 1 mM) for 1 hour at room temperature with shaking. Next, alkylation buffer (550 mM iodoacetamide/chloroacetamide in 50 mM ammonium bicarbonate) was added to the samples to a final concentration of 5.5 mM IAA (iodoacetamide), and samples were incubated for 1 hour at room temperature with shaking in the dark. LysC (2 µg/100 µg of protein) (NEB) was added, and the samples were incubated for 3 hours at room temperature, and then diluted with 4 volumes of 20 mM ammonium bicarbonate. Trypsin (2 µg/100 µg of protein) (Promega) was added, and the samples were incubated overnight at room temperature with shaking. After centrifugation (15,000 rpm, 10 minutes, room temperature), supernatants were transferred to new tubes, frozen at −80°C, and lyophilized. The digested samples were then analyzed by LC-MS/MS on an LTQ-Orbitrap instrument (Thermo).

#### YokF toxicity assays

##### Effect of YokF variants on cell growth

BSB1 (WT) or strains expressing YokF domains from an ectopic locus under an inducible promoter, were streaked over LB agar plates with or without IPTG (1 mM). Plates were incubated overnight at 37°C, and documented.

##### Growth dynamics

Overnight cultures of BSB1 (WT) or strains expressing YokF domains from an ectopic locus under an inducible promoter were diluted to OD_600nm_ 0.05 in the absence of IPTG (0.5 mM) and incubated at 37°C for 2 hours. Next, cultures were re-diluted (OD_600nm_ 0.05) into fresh LB with or without IPTG (0.5 mM), and growth kinetics were followed by measuring OD_600nm_ at 15 minute intervals.

##### Chromosome degradation assay

BSB1 (WT) or strains expressing YokF domains from an ectopic locus under an inducible promoter, were grown on LB agar plates with or without IPTG (1 mM), and harvested after 8 hours of incubation. Genomic DNA was extracted using Promega commercial kit, and DNA concentration was quantified using a NanoDrop spectrophotometer. For gel electrophoresis analysis, 200 ng of each DNA sample was subjected to agarose (1%) gel electrophoresis for 40 minutes at 100V. Gels were subsequently stained with ethidium bromide and visualized using a GelDoc Imaging System (Bio-Rad).

#### Whole genome sequencing and comparative analysis

Genomic DNA from PY79 and BSB1 *B. subtilis* strains was extracted using a commercially available kit (Promega). Single-end high-throughput sequencing (150 bp) was conducted using an Illumina NextSeq 500. Raw sequencing reads were processed to remove low-quality and technical sequences. Reads were trimmed using Cutadapt (version 1.15) to remove low-quality bases from both ends, retaining only reads of at least 30 bp in length after trimming. The trimmed reads were further filtered by overall quality using the fastq quality filter (FASTX package, version 0.0.14). The remaining reads were de-duplicated by exact sequence using an in-house Perl script. Processed reads were further analyzed using Geneious (version 11.1.5). Reads were aligned, the contigs were annotated to the *B. subtilis* PY79 reference genome (NCBI Accession no.: CP006881), and variations were manually examined. The raw sequence data for each strain were deposited in DDBJ/EMBL/GenBank under the project number PRJNA1098594 [SRA Accession: SRX24206248 (PY79), and SRX24206250 (BSB1)].

#### Structural prediction and computational analysis of YokF

The protein sequence of YokF was retrieved from the SubtiWiki^69^. Three-dimensional structural prediction was performed using AlphaFold protein structure database^50^, cross-referenced with its UniProt ID (AF-O32001-F1-v4). Signal peptide identification was performed using the SignalP 6.0 server^70^, while the catalytic and DNA binding sites were identified through HHPred^51^. Functional domain analysis was conducted using InterPro database^71^.

#### Phylogenetic analysis

YokF protein homologs containing all three domains were identified using BLASTP against the non-redundant protein database. The resulting dataset was manually curated to eliminate duplicates and ensure taxonomic diversity. Sequences were aligned using ClustalW method. A phylogenetic tree of YokF homologs was constructed using the neighbor-joining algorithm in MEGA 7.0, with 1000 bootstrap replicates, and visualized using the online platform TVBOT v2.6. Multiple sequence alignment was performed using DNAMAN sequence analysis software.

Next, YokF homologs were manually curated using BLASTP (database: non-redundant protein sequences), and their genetic context was analyzed. A 20 kb region on either side of each homolog was examined for the presence of phage-encoded proteins. Homologs were classified as “putatively host-encoded”, when no phage-encoded proteins were identified within the 40 kb region. Additionally, the homologs were cross-referenced with the Prokaryotic Virus Remote Homologous Groups (PHROGs) database^72^ (https://phrogs.lmge.uca.fr/) to detect any phage-related sequences.

For the functional analysis, nucleotide sequences of selected YokF homologs were retrieved using TBLASTN (database: core Nucleotide). The corresponding DNA sequences were commercially synthesized (Twist Bioscience) and engineered into *B. subtilis* (PY79) under an IPTG-inducible promoter and assessed for their NPex inhibitory activity.

#### Quantification and statistical analysis

GraphPad Prism was used for all statistical analyses, data processing, and presentation. Unless stated otherwise, bar charts and graphs display a mean ± SEM from at least 3 independent repeats. All quantifications of microscopy images were performed with several technical and biological replicates, as indicated in the respective figure legends.

---

**Fig. S1.**
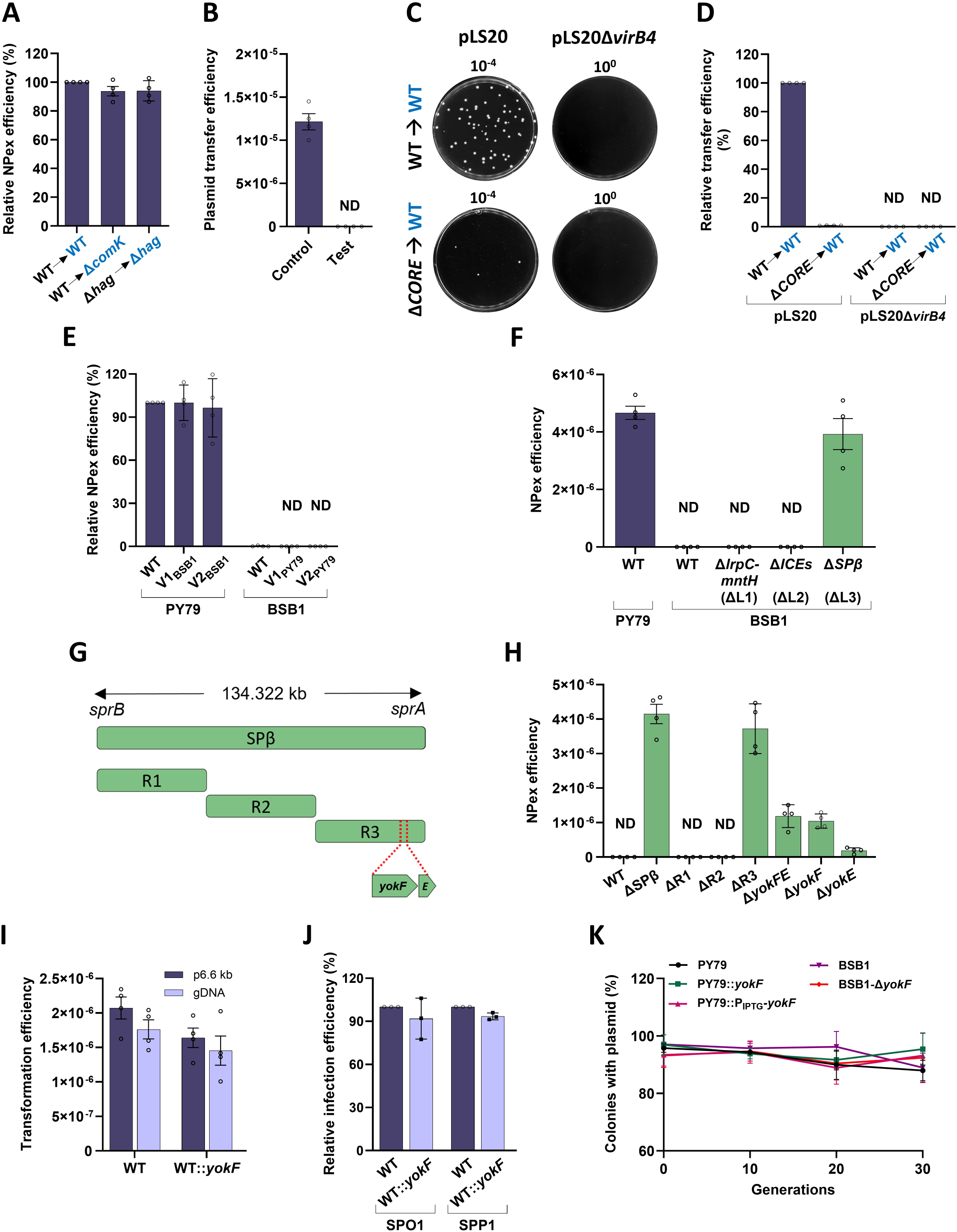
NPex is inhibited by YokF. **(A)** <u>NPex is competence and motility independent</u>: PY79-derived donor strains: WT (PY79) (GV478), and Δ*hag* (GB01: Δ*hag*::*erm*), carrying p6.6 kb (pHB201/*cat*, *erm*), were mixed in a 1:1 ratio with kanamycin resistance recipient strains: WT (SB513: *amyE*::P_IPTG_-*gfp*-*kan*), Δ*comK* (GB252: Δ*comK*::*tet*, *amyE*::P_IPTG_-*gfp*-*kan*), and Δ*hag* (GD268: Δ*hag*::*erm*, *amyE*::P_IPTG_-*gfp*-*kan*). The used donor (black) and recipient (blue) pairs are indicated. The mixtures were incubated for 4 hours without selection, and equal volumes of cells were then spread over plates containing chloramphenicol, erythromycin, and kanamycin to select for trans-recipients. NPex efficiency was determined as the ratio of trans-recipients (CFU)/total recipients (CFU). Shown is NPex efficiency (%) relative to WT → WT pair. Data is presented as mean values and SEM, based on 4 independent experiments. **(B)** <u>NPex is transduction- and membrane vesicle-independent</u>: Wild-type PY79-derived donor (GV478) and kanamycin resistance recipient (SB513: *amyE*::P_IPTG_*-gfp-kan*) were mixed in 1:1 ratio and incubated for 4 hours without selection. Equal volumes of cells were then spread over plates containing chloramphenicol, erythromycin, and kanamycin to select for trans-recipients (Control). In parallel, donor and recipient cells were cultured separately for 3 hours without selection. The spent medium of the recipient strain was then replaced with filtered supernatant from the donor strain culture, followed by incubation for 1 hour. Equal volumes of cells were then spread over plates containing chloramphenicol, erythromycin, and kanamycin to select for trans-recipients (Test). Plasmid transfer efficiency was determined as the ratio of trans-recipients (CFU)/total recipients (CFU). Data is presented as mean values and SEM, based on 4 independent experiments. **(C)** <u>NPex is limited by plasmid size</u>: PY79-derived donor strains: WT (PY79), and Δ*CORE* (Δ*fliO-flhA*::*tet,* P*_fla/che_*-*flhF*) carrying pLS20/*cm* (SH337 and SH352, respectively) or pLS20Δ*virB4*/*cm* (SH346 and GV666, respectively) were mixed in 1:1 ratio with kanamycin resistance WT recipient strain (SB513: *amyE*::P_IPTG_-*gfp*-*kan*). The used donor (black) and recipient (blue) pairs are indicated. The mixtures were incubated for 4 hours without selection, and equal volumes of cells were then spread over plates containing chloramphenicol, and kanamycin to select for recipients that acquired the plasmids. Shown are representative images of selective plates at the indicated dilutions. Of note, *CORE* was found to be involved in facilitating pLS20 conjugation, hence resulting in low conjugation efficiency of the mutant^73^. **(D)** Quantitative analysis of plasmid transfer efficiencies between donor and recipient pairs shown in (B). Plasmid transfer efficiency was determined as in (A). Shown is plasmid transfer efficiency (%) relative to WT/pLS20 → WT pair. Data are presented as mean values and SEM, based on 4 independent experiments. ND – Not Detected. **(E)** <u>Mapping NPex inhibition genomic locus</u>: Donor strains: WT (PY79) (GV478), and PY79-derived strains harboring V1_BSB1_(GV669), or V2_BSB1_ (GV671), and WT (BSB1) (GV345), and BSB1-derived strains harboring V1_PY79_ (GV377), or V2_PY79_ (GV384), all carrying p6.6 kb (pHB201/*cat*, *erm*), were mixed in a 1:1 ratio with kanamycin resistance recipient strains (SB513, GV670, GV672, GV372, GV375, and GV382), harboring equivalent genotypes, but lacking a plasmid. The mixtures were incubated for 4 hours without selection, and equal volumes of cells were then spread over plates selective for trans-recipients. A detailed description of the used donor and recipient pairs and antibiotic selections employed are listed in Table S1. NPex efficiency was calculated as in (A). Shown is NPex efficiency (%) relative to WT → WT (PY79) pair. Data are presented as mean values and SEM, based on 4 independent experiments. ND – Not Detected. **(F)** <u>SPβ prophage inhibits NPex</u>: Donor strains: WT (PY79) (GV478), WT (BSB1) (GV345) and BSB1-derived strains, ΔL1 (GV674: Δ*lrpC-mntH*::*tet*), ΔL2 (SH624: Δ*ydcL*-*yddM*::*tet*), and ΔL3 (SH625: Δ*sprB*-*sprA*::*tet*), all carrying p6.6 kb (pHB201/*cat*, *erm*), were mixed in a 1:1 ratio with kanamycin resistance recipient strains (*amyE*::P_IPTG_*-gfp-kan*) harboring equivalent genotypes (SB513, GV372, GV675, SH620, and SH621), but lacking a plasmid. The mixtures were incubated for 4 hours without selection, and equal volumes of cells were then spread over plates containing chloramphenicol, erythromycin, and kanamycin to select for trans-recipients. NPex efficiency was calculated as described in (A). Data are presented as mean values and SEM, based on 4 independent experiments. **(G)** <u>Sequential deletion analysis of SPβ genome</u>: Schematic illustration of the sequential deletion analysis of SPβ prophage genome in BSB1 for identifying the NPex inhibitory region. SPβ was divided into 3 regions (R1: *sprB*-*yopS*, R2: *yopP*-*youB*, and R3: *yomK*-*sprA*). *YokFE* operon is located in R3. **(H)** <u>Sequential deletion analysis</u>: BSB1-derived donor strains: WT (BSB1) (GV345), ΔSPβ (SH625), ΔR1 (GV119), ΔR2 (GV239), ΔR3 (SH627), Δ*yokFE* (GV428), Δ*yokF* (GV417), and Δ*yokE* (GV419), all carrying p6.6 kb (pHB201/*cat*, *erm*), were mixed in a 1:1 ratio with kanamycin resistance recipient strains: (GV372, SH621, GV118, GV238, SH623, GV427, GV416, and GV418), harboring equivalent genotypes but lacking a plasmid. The mixtures were incubated for 4 hours without selection, and equal volumes of cells were then spread over plates selective for trans-recipients. R1-R3 regions correspond to (F). A detailed description of the used donor and recipient pairs and antibiotic selections employed are listed in Table S1. NPex efficiency was calculated as in (A). Data are presented as mean values and SEM, based on 4 independent experiments. ND – Not Detected. **(I)** <u>Natural transformation is unaffected by YokF</u>: WT (PY79) or PY79 expressing YokF, WT::*yokF* (GV462: *sacA*::P*_yokF_*-*yokF*_BSB1_-*spec*) cells were grown under competence-inducing conditions with either genomic DNA (100 ng) encoding a tetracycline resistance gene, or p6.6 kb (pHB201/*cat*, *erm*) (100 ng). An equal volume of cells was spread over plates containing tetracycline or chloramphenicol to select for transformants, and on antibiotic-free media for the total population. Transformation efficiency was determined as the ratio of transformants (CFU)/total population (CFU). Data are presented as mean values and SEM, based on 4 independent experiments. **(J)** <u>YokF does not affect phage sensitivity</u>: WT (PY79) or PY79 expressing YokF, WT::*yokF* (GV462: *sacA*::P*_yokF_*-*yokF*_BSB1_-*spec*) cells were grown to mid-logarithmic phase and infected with SPO1 or SPP1 phages. After phage attachment, infected cells were plated onto MB agar plates. Infection efficiency was determined as the ratio of plaque forming units (PFU) between WT::*yokF* and WT. Shown is relative infection efficiency (%). Data are presented as mean values and SEM, based on 3 independent experiments. **(K)** <u>Plasmid stability is unaffected by YokF:</u> WT (PY79) (GV478), PY79 expressing YokF PY79::*yokF* (GV469: *sacA*:: P*_yokF_*-*yokF*_BSB1_-*spec*), PY79::P_IPTG_*-yokF* (GV480: *amyE*::P_IPTG_*-yokF*_BSB1_-*spec*), WT (BSB1) (GV345), and isogenic Δ*yokF* (GV417: Δ*yokF*::*tet*) strains, all carrying p6.6 kb (pHB201/*cat*, *erm*), were grown with antibiotics (generation 0). PY79::P_IPTG_*-yokF* was grown in the presence of IPTG. Cells were refreshed every 3 hours (∼10 generations) without antibiotics until 30 generations. Every 10 generations, equal volumes of cells were spread over plates containing chloramphenicol and antibiotic-free media to estimate the numbers of plasmid-containing cells and the total population, respectively. The graph illustrates plasmid stability as the ratio of plasmid-containing cells (CFU)/total population (CFU). Data is presented as mean values and SEM, based on 4 independent experiments.

**Fig. S2.**
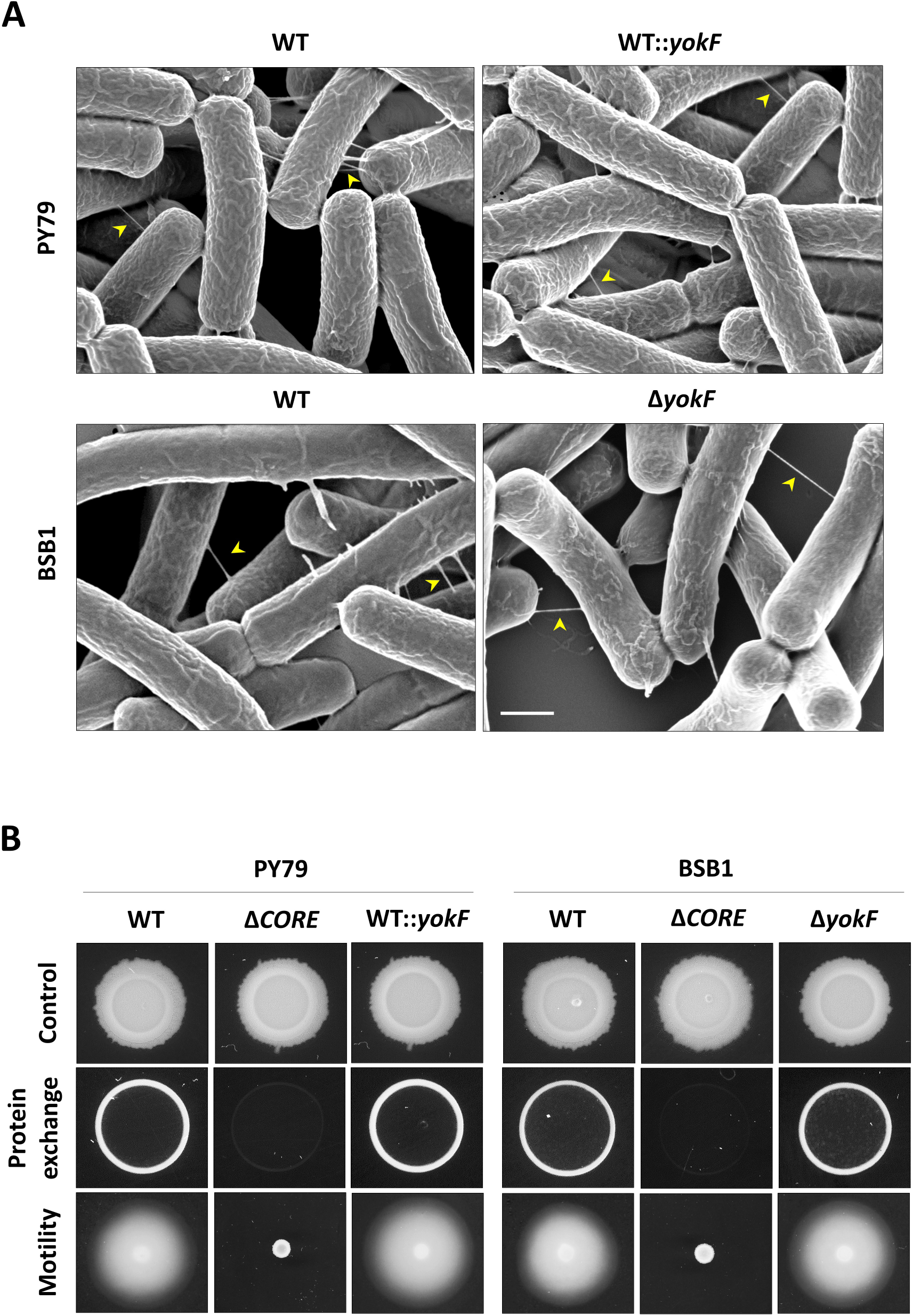
YokF does not inhibit nanotube formation or protein exchange. **(A)** <u>Nanotube formation appears unaffected by YokF</u>: WT (PY79), PY79 expressing YokF PY79::*yokF* (GV462: *sacA*:: P*_yokF_*-*yokF*_BSB1_-*spec*), WT (BSB1) (GV345), and isogenic Δ*yokF* (GV411: Δ*yokF*::*tet*) were grown to the mid-logarithmic phase, spotted onto EM grids, incubated on LB agar plates for 4 hours at 37°C, and visualized by HR-SEM. Arrows indicate intercellular nanotubes connecting neighboring cells. Scale bar 0.5 µm. **(B)** <u>Protein exchange is unaffected by YokF</u>: WT (PY79) (SB463: *amyE*::P_IPTG_*-cat-spec*), isogenic Δ*CORE* (SH17: Δ*fliO-flhA*::*tet,* P*_fla/che_-flhF*, *amyE*::P_IPTG_*-cat-spec*), WT::*yokF* (GV274: *sacA*::*yokF*_BsB1_-*spec*, *amyE*::P_IPTG_*-cat-spec*), WT (BSB1) (GV373: *amyE*::P_IPTG_*-cat-spec*), isogenic Δ*CORE* (GV494: Δ*fliO-flhA*::*tet,* P*_fla/che_-flhF*, *amyE*::P_IPTG_*-cat-spec*), and isogenic Δ*yokF* (GV456: Δ*yokF*::*tet, amyE*::P_IPTG_*-cat-spec*) were mixed in a 1:1 ratio with kanamycin resistance strains (*amyE*::P_IPTG_*-gfp-kan*) harboring equivalent genotypes (SB513, SH13, GV468, GV372, GV107, and GV416). The mixtures were incubated for 4 hours without selection, and equal volumes of cells were spotted onto LB agar (control) and LB agar with chloramphenicol and kanamycin to evaluate protein exchange. Plates were photographed after 18 hours of incubation. For motility assay, PY79 (WT), isogenic Δ*CORE* (SH9: Δ*fliO-flhA*::*tet,* P*_fla/che_-flhF*), WT::*yokF* (GV462: *sacA*::*yokF*_BsB1_-*spec*), BSB1(WT), isogenic Δ*CORE* (GV582: Δ*fliO-flhA*::*tet,* P*_fla/che_-flhF*), and isogenic Δ*yokF* (GV411: Δ*yokF*::*tet*) strains were grown to the mid-logarithmic phase and spotted onto 0.3% agar LB plates. Plates were incubated for 7 hours at 37°C before being photographed.

**Fig. S3.**
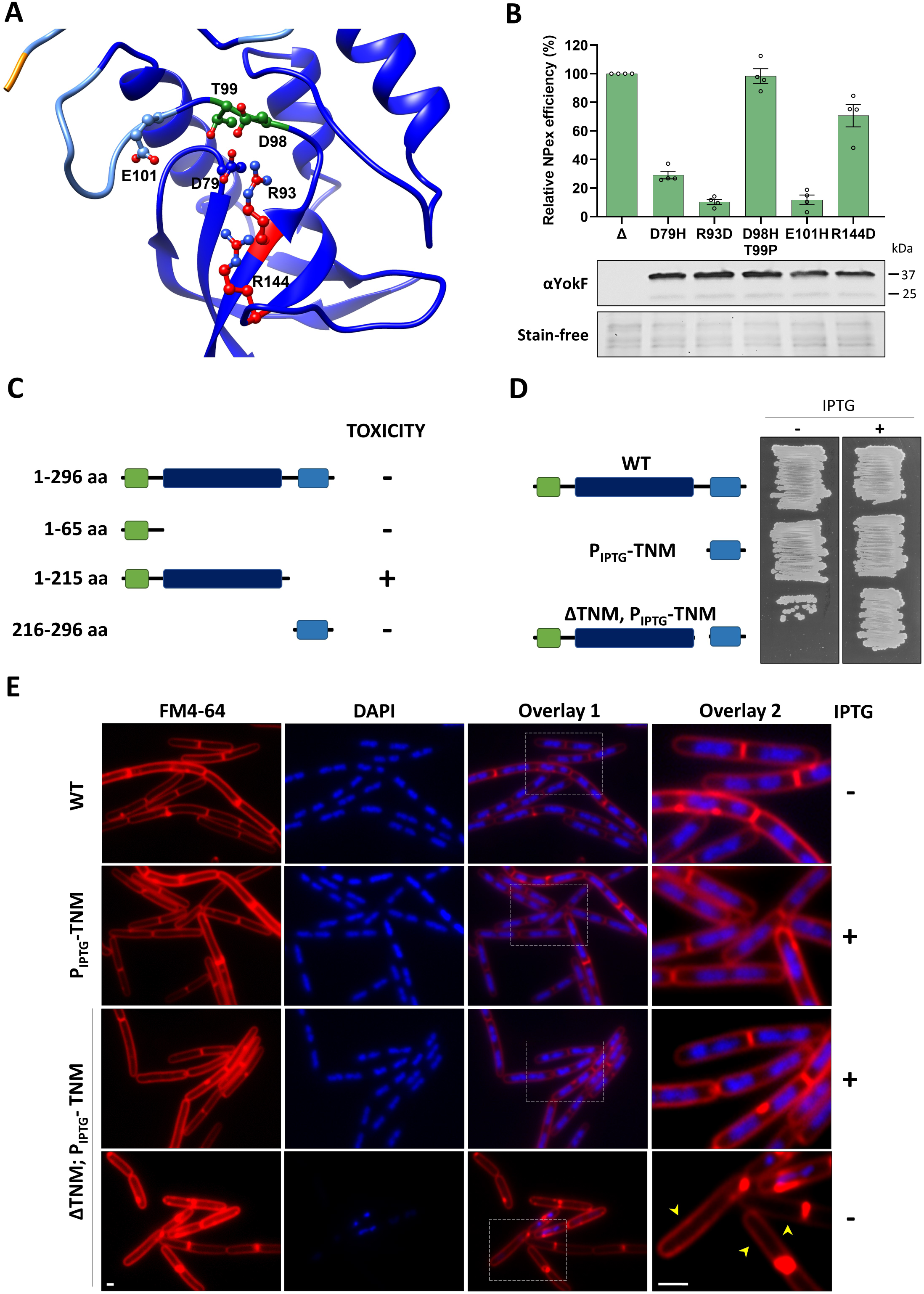
YokF nuclease activity is modulated and essential for NPex inhibition. **(A)** <u>YokF TNase domain structure</u>: AlphaFold-predicted structure of YokF TNase domain, emphasizing the potential active and binding site residues D79, R93, D98, T99, E101, and R144 clustered within the TNase domain. High pLDDT (predicted local difference test) value is shown in a blue ribbon, indicating a high level of confidence in the structural prediction. **(B)** <u>YokF TNase mutants are NPex-deficient</u>: Upper panel: BSB1-derived donor strains: Δ*yokF* (GV417), *yokF*_D79H_ (GV602), *yokF*_R93D_ (GV576), *yokF*_D98H-T99P_ (GV604), *yokF*_E101H_ (GV578), and *yokF*_R144D_ (GV580), carrying p6.6 kb (pHB201/*cat*, *erm*), were mixed in a 1:1 ratio with kanamycin resistance recipient strains (*amyE*::P_IPTG_*-gfp-kan*) (GV416, GV601, GV575, GV603, GV577, and GV579), harboring equivalent genotypes, but lacking a plasmid. The mixtures were incubated for 4 hours without selection, and equal volumes of cells were then spread over plates selective for trans-recipients. A detailed description of the used donor and recipient pairs and antibiotic selections employed are listed in Table S1. NPex efficiency was determined as the ratio of trans-recipients (CFU)/total recipients (CFU). Shown is NPex efficiency (%) relative to Δ*yokF* pair. Data is presented as mean values and SEM, based on 4 independent experiments. Lower panels: BSB1-derived strains expressing the indicated YokF mutants (GV411, GV598, GV566, GV599, GV567, and GV568) were grown to mid-logarithmic phase and the total lysates were extracted. Shown is a western blot analysis using anti-YokF antibodies, with a stain-free image as a loading control. **(C)** Schematic illustration of YokF truncations and their associated toxicity when expressed from the native chromosomal locus in BSB1. Toxicity (+) was manifested by the failure to obtain colonies following transformation, whereas lack of toxicity (-) showed no impact on transformation efficiency. **(D)** <u>YokF exhibits a self-inhibiting TNM domain</u>: BSB1-derived cells expressing YokF (WT), P_IPTG_-TNM (GV573: *amyE*::P_IPTG_*-yokF*_216-296aa_-*spec*), and ΔTNM, P_IPTG_-TNM (GV619: *yokF*_1-215aa_-*tet*, *amyE::*P_IPTG_-*yokF*_216-296aa_-*spec*), were streaked on LB agar plates with (+) or without (-) IPTG as indicated, and incubated overnight at 37°C. Shown are representative images of at least 3 independent experiments. The residual growth for ΔTNM, P_IPTG_-TNM without IPTG was found to emanate from suppressor mutations in the *yokF* region (data not shown). **(E)** <u>Uninhibited TNase domain causes chromosome degradation</u>: BSB1 cells expressing YokF (WT), P_IPTG_-TNM (GV573: *amyE*::P_IPTG_*-yokF*_216-296aa_-*spec*), and ΔTNM, P_IPTG_-TNM (GV619: *yokF*_1-215aa_-*tet*, *amyE::*P_IPTG_-*yokF*_216-296aa_-*spec*), were grown for 2 hours with (+) or without (-) IPTG, as indicated. Shown are FM4-64 (red), DAPI (blue) fluorescence images, their overlay (overlay 1), and the enlargement of the inset (overlay2). Arrows highlight cells lacking a signal from DAPI staining. Scale bars 0.5 µm.

**Fig. S4.**
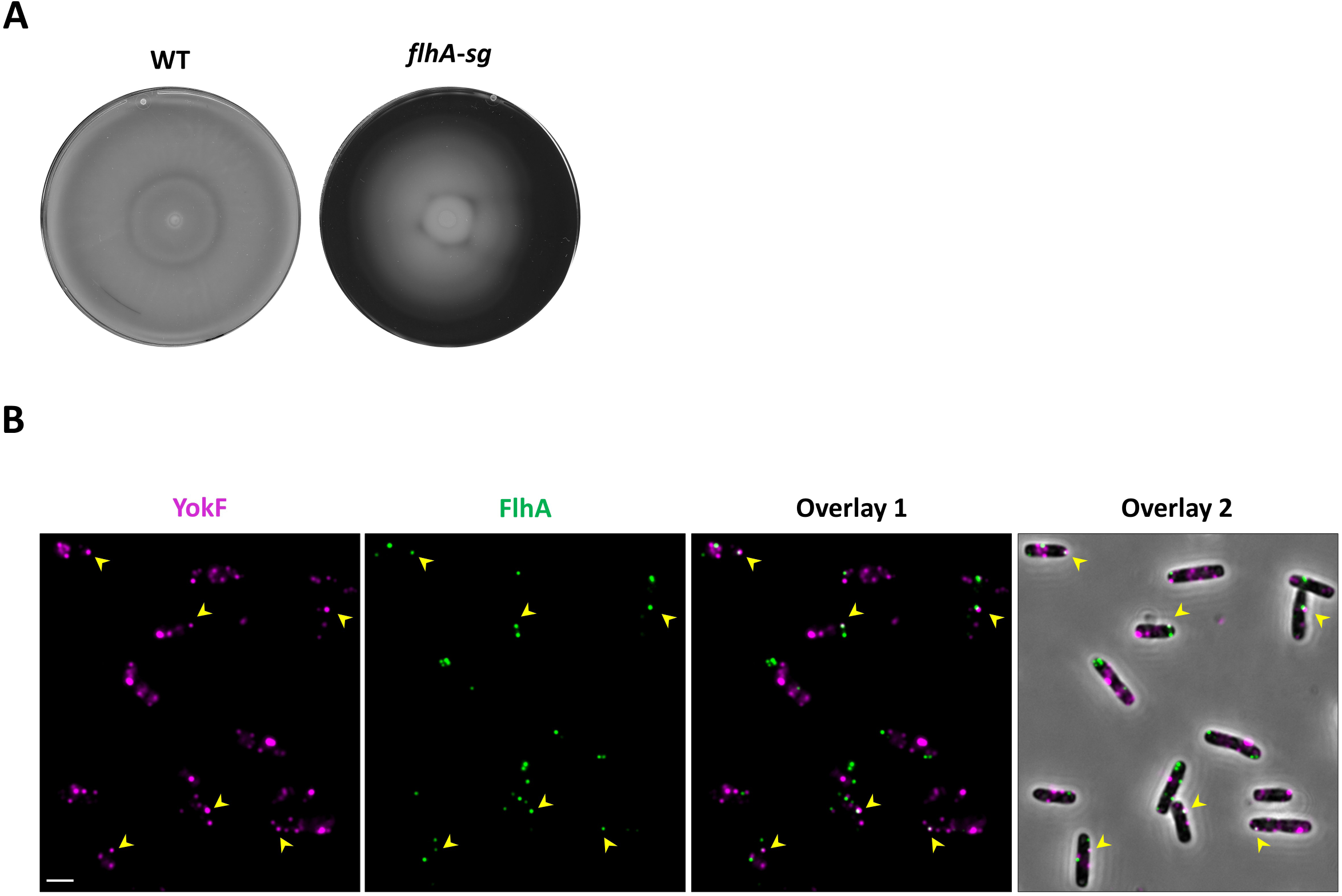
YokF colocalizes with FlhA. **(A)** *<u>flhA-sg</u>* <u>is motile</u>: BSB1 (WT), and *flhA*-*sg* (GV618: *flhA-lin_4x_-sg-Kan-*P*_fla/che_-flhF*) strains were grown to the mid-logarithmic phase, spotted onto 0.3% agar LB plates, and photographed after 12 hours of incubation at 37°C. **(B)** <u>YokF and FlhA colocalization analysis</u>: BSB1-derived strain expressing FlhA-SG (GV617: *flhA-lin_4_*_x_*-sg_d_-kan*-P*_fla/che_*-*flhF*) was treated with anti-YokF primary antibodies followed by Alexa 647-conjugated secondary antibodies and subjected to fluorescence microscopy. Shown are fluorescence images of YokF (magenta), FlhA-SG (green), their overlay (Overlay 1), and their overlay with phase contrast (gray) (Overlay 2). Arrows highlight YokF and FlhA colocalization sites. Scale bar 1 µm.

**Fig. S5.**
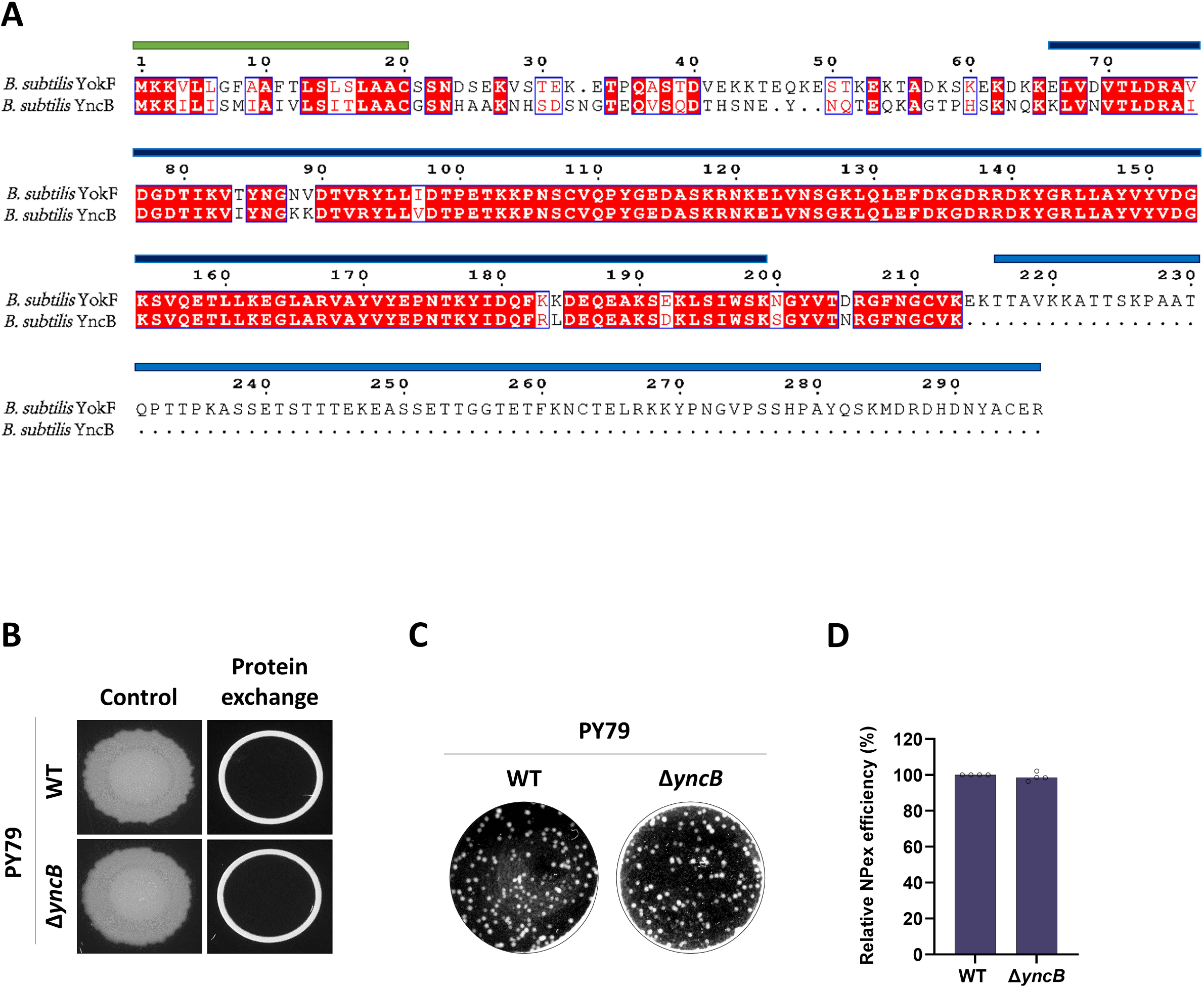
Comparing YokF with its putative paralogue YncB. **(A)** <u>YncB-YokF sequence comparison</u>: Comparative sequence alignment of YokF and YncB from *B. subtilis*. Protein sequences were aligned using T-COFFEE and annotated with ESPript. Similar and identical amino acid residues are shown in red, highlighted within blue frames. Domain boundaries are delineated by colored bars: SP (green), TNase (dark blue), and TNM (light blue). **(B)** <u>YncB does not impact nanotube-mediated protein exchange</u>: PY79-derived strains WT (SB463: *amyE*::P_IPTG_*-cat-spe*c), and Δ*yncB* (GV473: Δ*yncB*::*tet, amyE*::P_IPTG_*-gfp-kan*), were mixed in a 1:1 ratio with kanamycin resistance recipients strains (*amyE*::P_IPTG_*-gfp-kan*) harboring equivalent genotypes (SB513, and GV474). The mixtures were incubated for 4 hours without selection, and equal volumes of cells were spotted onto LB agar (control) and LB agar with chloramphenicol and kanamycin to evaluate protein exchange. Plates were photographed after 18 hours of incubation. **(C)** <u>YncB does not affect NPex</u>: PY79-derived donor strains: WT (GV478), and Δ*yncB* (GV472: Δ*yncB*::*tet*), carrying p6.6 kb (pHB201/*cat*, *erm*), were mixed in a 1:1 ratio with kanamycin resistance recipient strains (*amyE*::P_IPTG_-*gfp*-*kan*), harboring equivalent genotypes (SB513, and GV474). The mixtures were incubated for 4 hours without selection, and equal volumes of cells were then spread over plates containing chloramphenicol, erythromycin, and kanamycin to select for trans-recipients. Shown are representative images of NPex selective plates. **(D)** Quantitative analysis of NPex efficiencies between the donor and recipient pairs shown in (C). NPex efficiency=Trans-recipients CFU/Total recipients CFU. Shown is NPex efficiency (%) relative to the WT pair. Data are presented as mean values and SEM, based on 4 independent experiments.

**Fig. S6.**
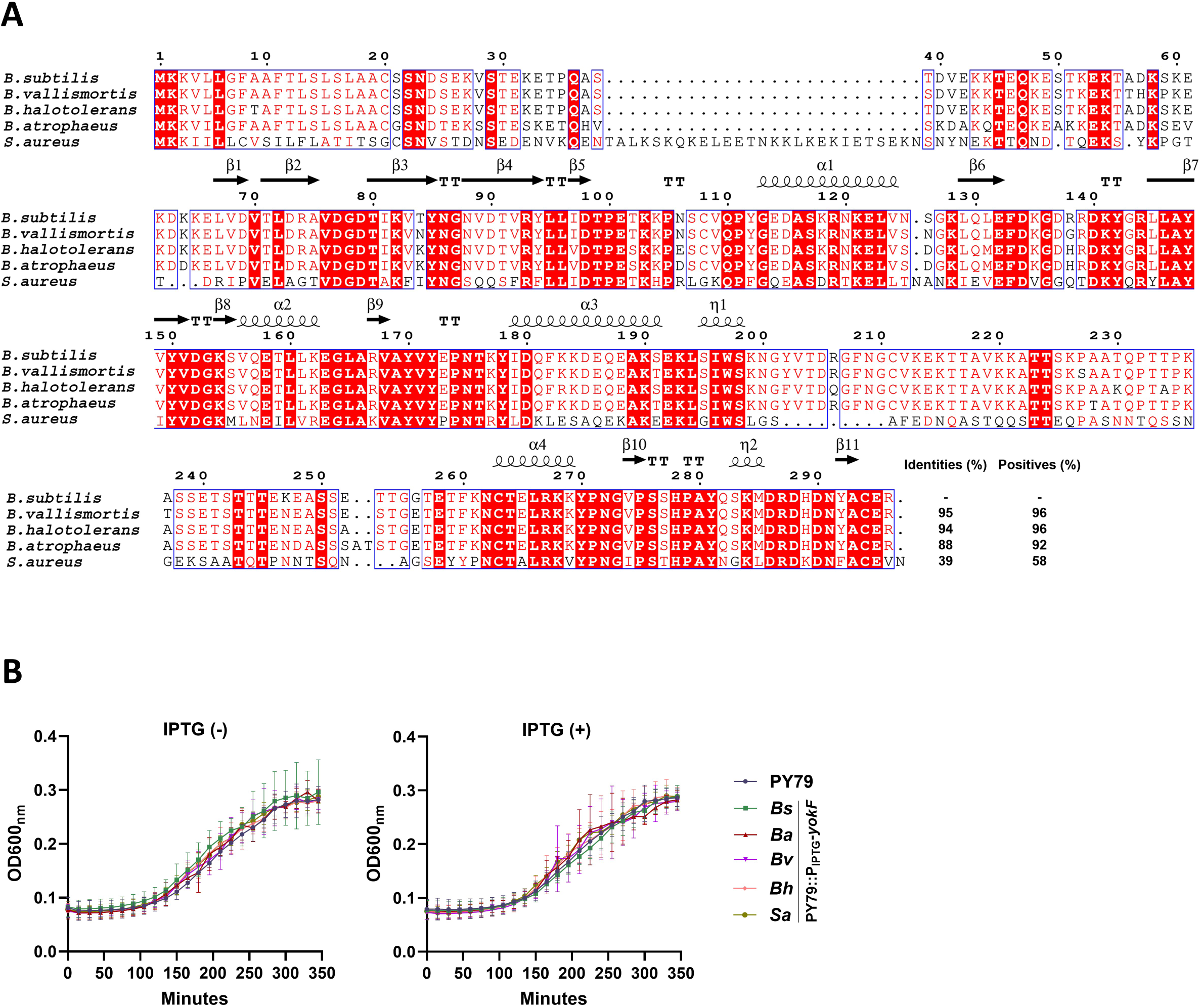
Characterization of YokF homologs. **(A)** <u>Multiple sequence alignment of YokF homologs</u>: Comparative sequence alignment of YokF from various bacterial species obtained from GenBank (Table S3). YokF homologs were aligned using T-COFFEE, and annotated with ESPript. Similar and identical amino acid residues are shown in red, highlighted within blue frames, with the percentage of identity and similarity between the YokF sequence from *B. subtilis* and its homologs indicated. Secondary structural elements derived from the *B. subtilis* YokF predicted structure are depicted above the sequences. **(B)** <u>Expression of YokF homologs has no growth impact</u>: PY79-derived cells expressing YokF homologs from *B. subtilis* (BSB1) (*Bs*) (GV480: *amyE*::P_IPTG_*-yokF_Bs_*-*spec*), *B. atrophaeus* (*Ba*) (GV653: *amyE*::P_IPTG_*-yokF_Ba_*-*spec*), *B. vallismortis* (*Bv*) (GV655: *amyE*::P_IPTG_*-yokF_Bv_*-*spec*), *B. halotolerans* (*Bh*) (GV657: *amyE*::P_IPTG_*-yokF_Bh_*-*spec*), and *S. aureus* (*Sa*) (GV659: *amyE*::P_IPTG_*-yokF_Sa_*-*spec*) were grown with (+) or without (-) IPTG. Cell growth was followed by measuring OD_600nm_ at 15 minute intervals. Data is presented as mean values and SEM, based on at least 3 independent repeats.

**Table S1.** A list of donor-recipient strain pairs and selective antibiotics used for NPex assays, as referenced in Fig. 1A, 1G, S1D, S1G, and S3B.

**Table S2.** Mass spectrometry (MS) analysis of the YokF interactome using PY79-derived strains: GV549 (Δ*yncB*::*tet*, Δ*hag*::*mls*) (Control), and GV556 (*sacA*::*yokF*_BSB1_-*spec*, Δ*yncB*::*tet*, Δ*hag*::*mls*). Following immunoprecipitation (IP) with anti-YokF antibodies, the co-immunoprecipitated proteins were analyzed by MS. The displayed values indicate the abundance of these proteins as determined by MS. Data shown are from a representative experiment of 2 biological repeats.

**Table S3.** A list of bacterial species harboring **YokF homologs** and their **genetic context**, related to Fig. 4A-4C, and S6A-S6B. YokF homologs, containing all three domains (SP, TNase, and TNM) were identified using BLASTP and curated to remove duplicates. The flanking regions of each 44 selected YokF homolog were examined for any phage-encoded proteins, and the genetic context was classified as ‘Host’ if no phage proteins were detected within the ∼40 kb region. Listed are NCBI accession IDs, YokF-amino acid sequences for each strain and the whole genome sequence (WGS) accession IDs used for the analysis. Red stars indicate homologs tested for functionality. N/A – Not Applicable.

**Table S4.**
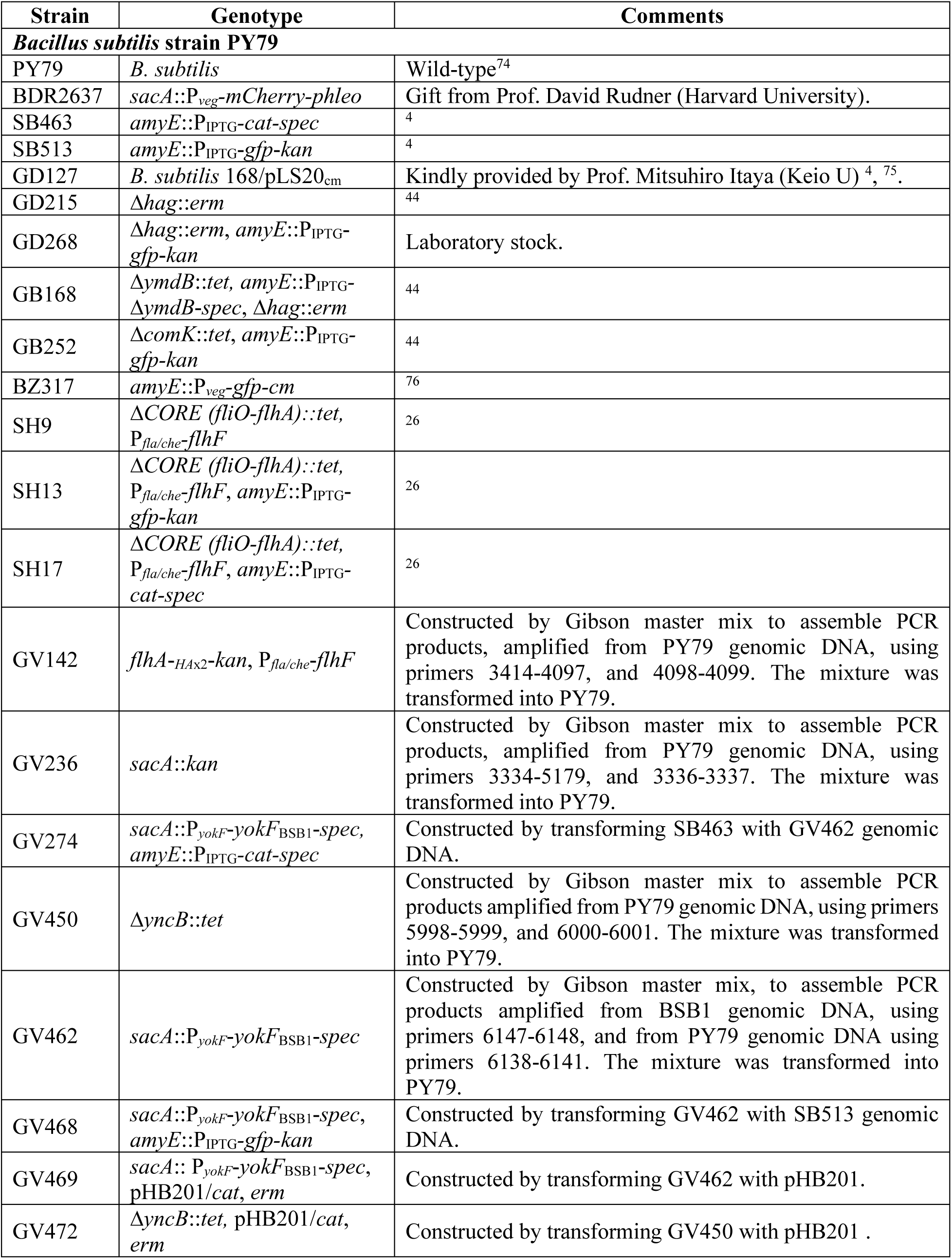
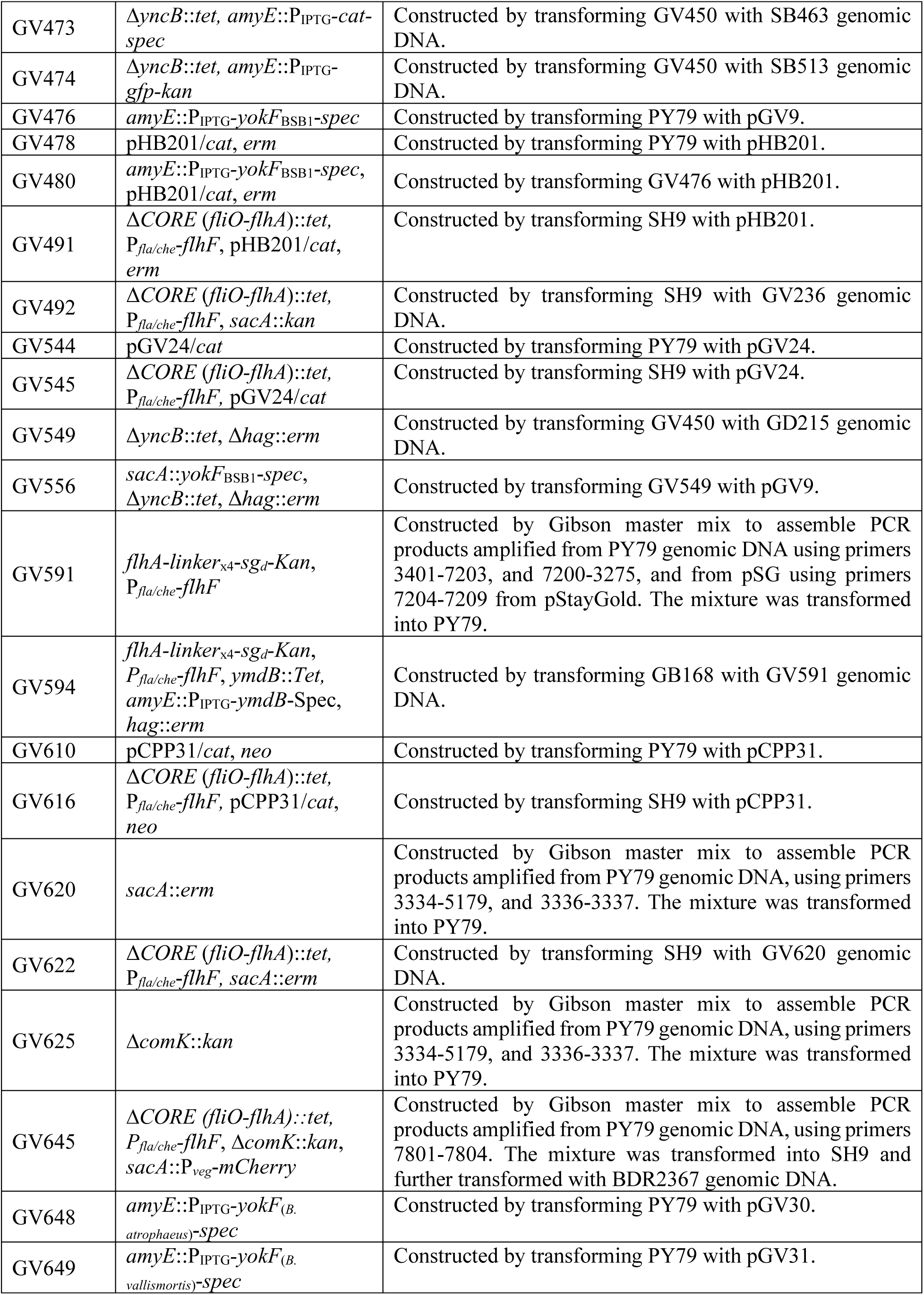
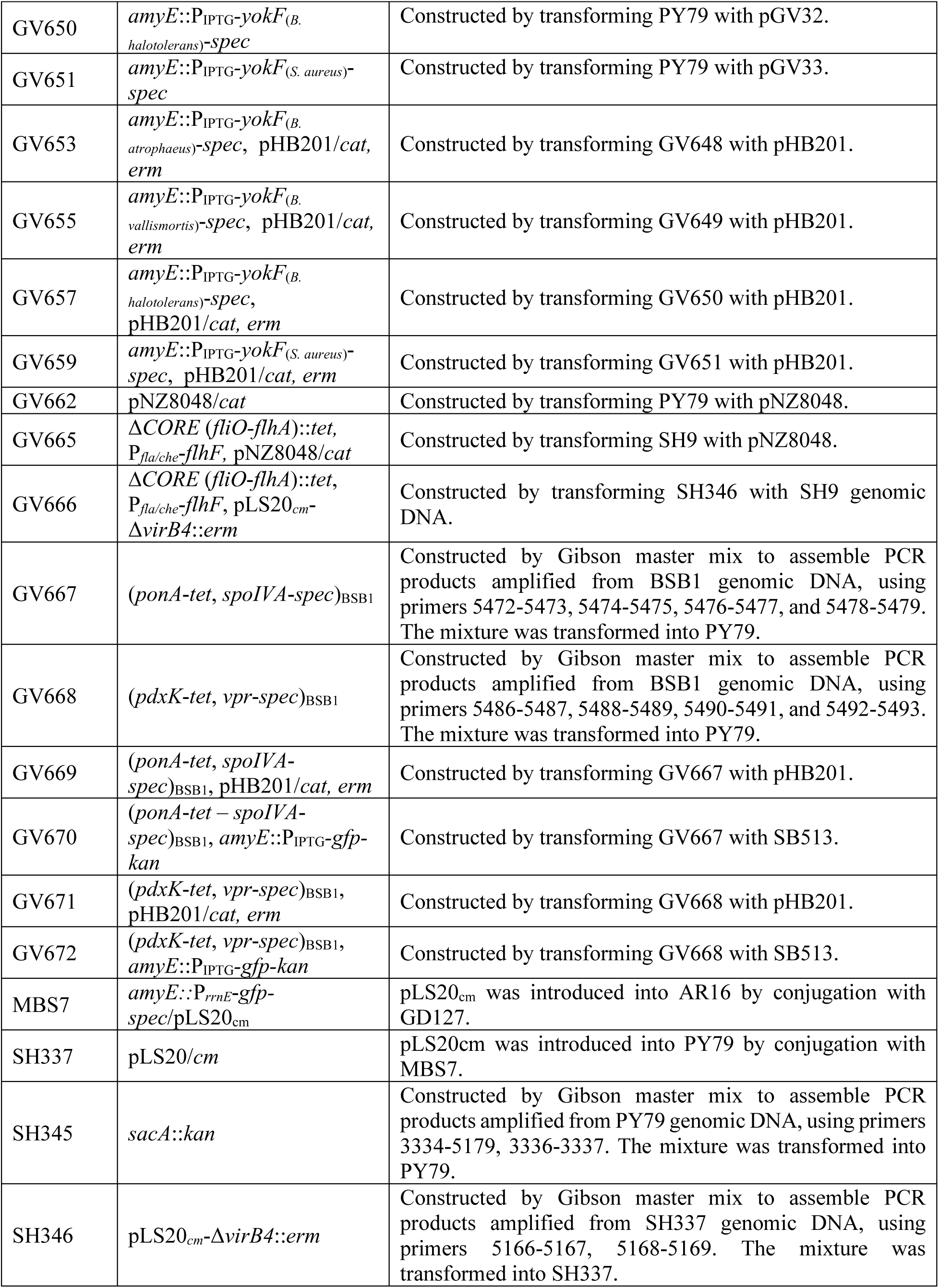
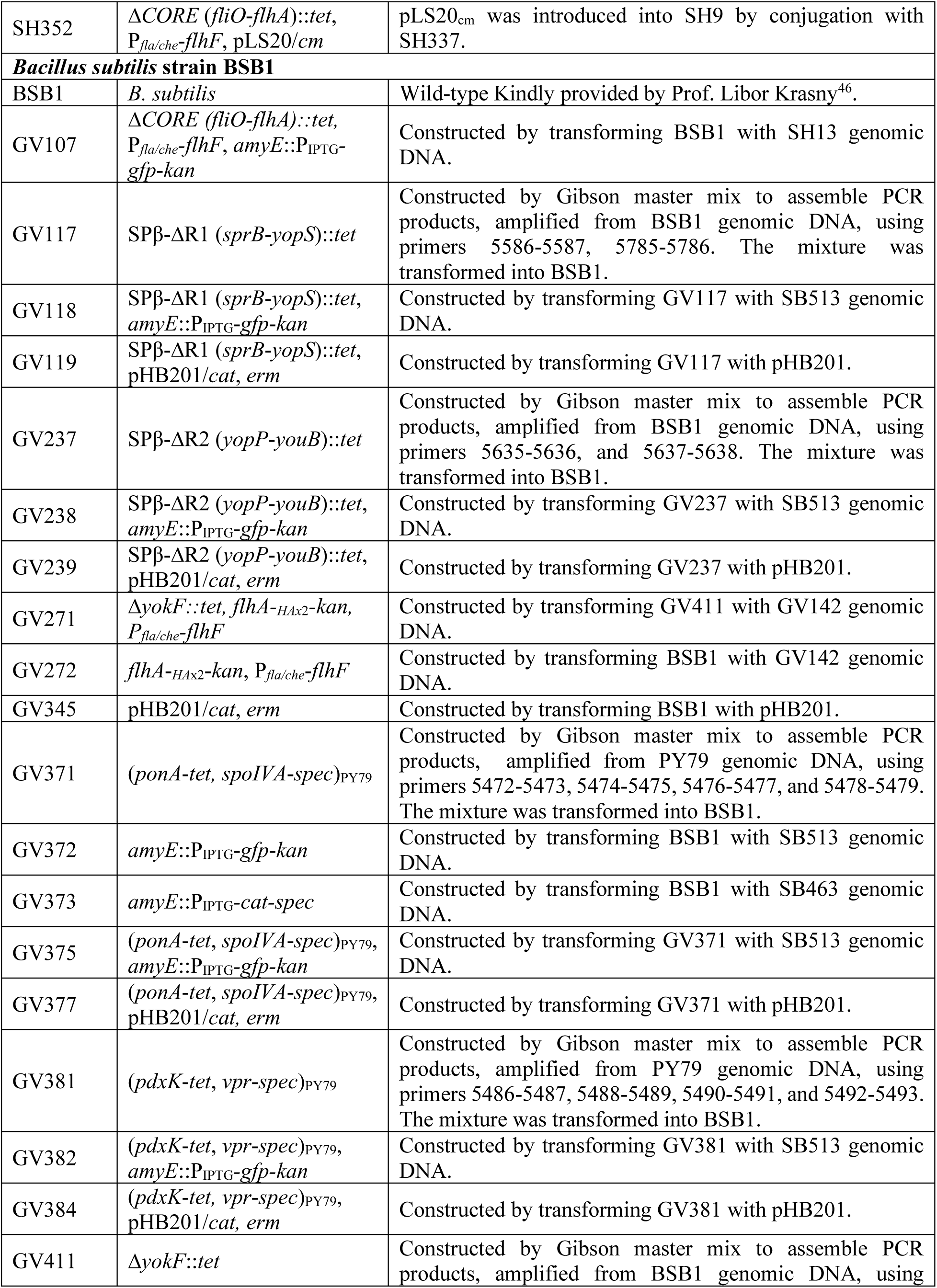
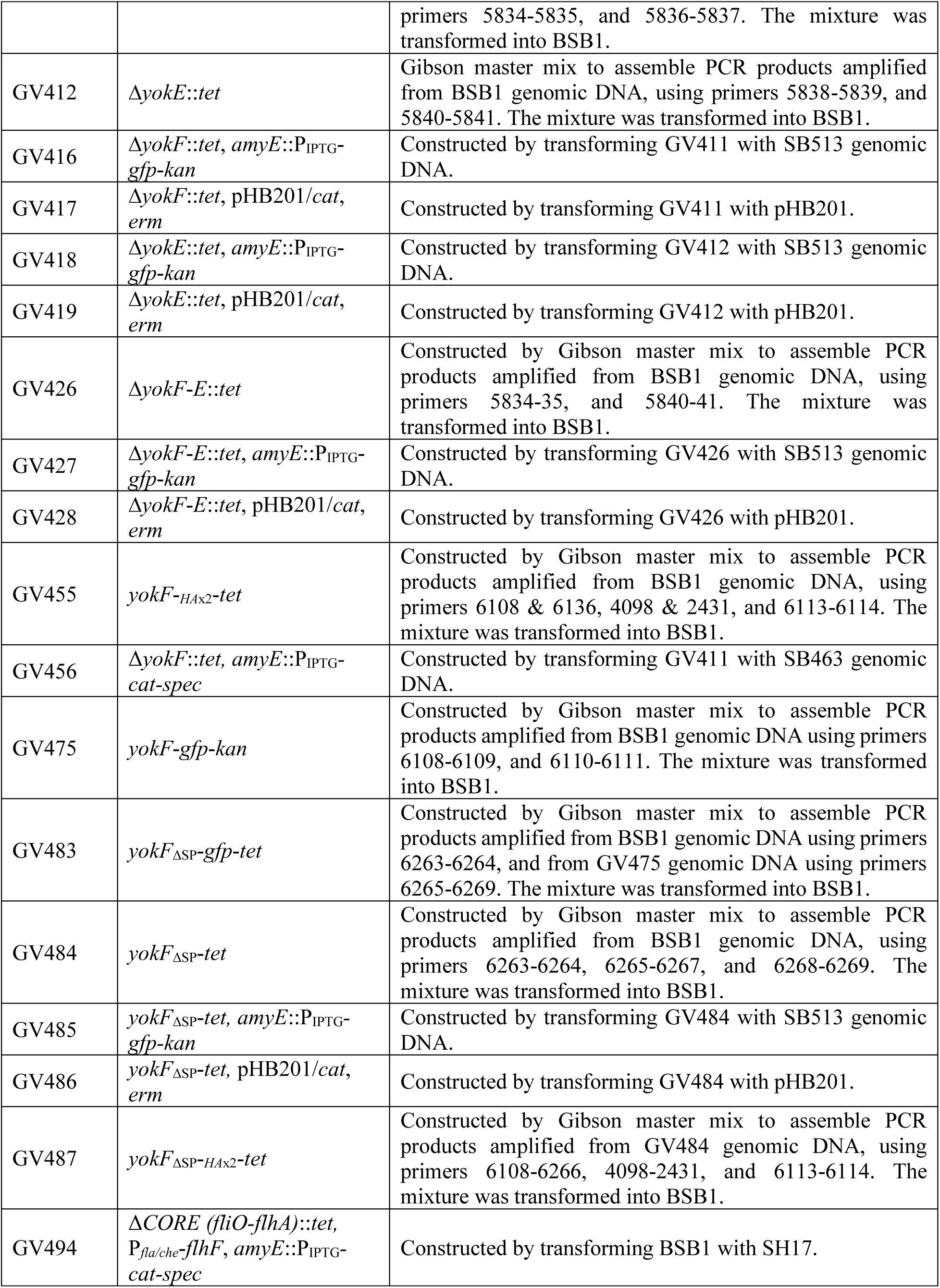
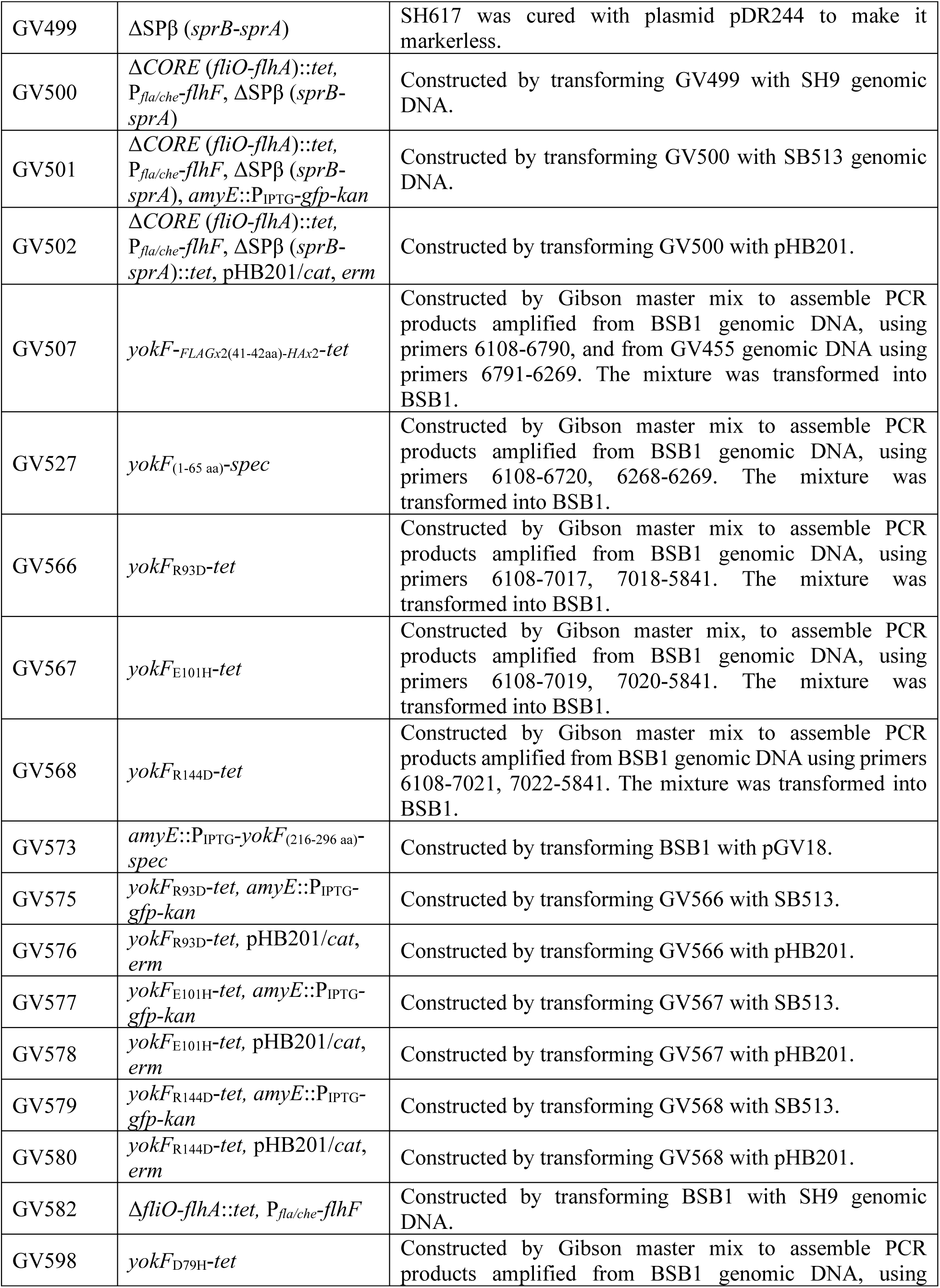
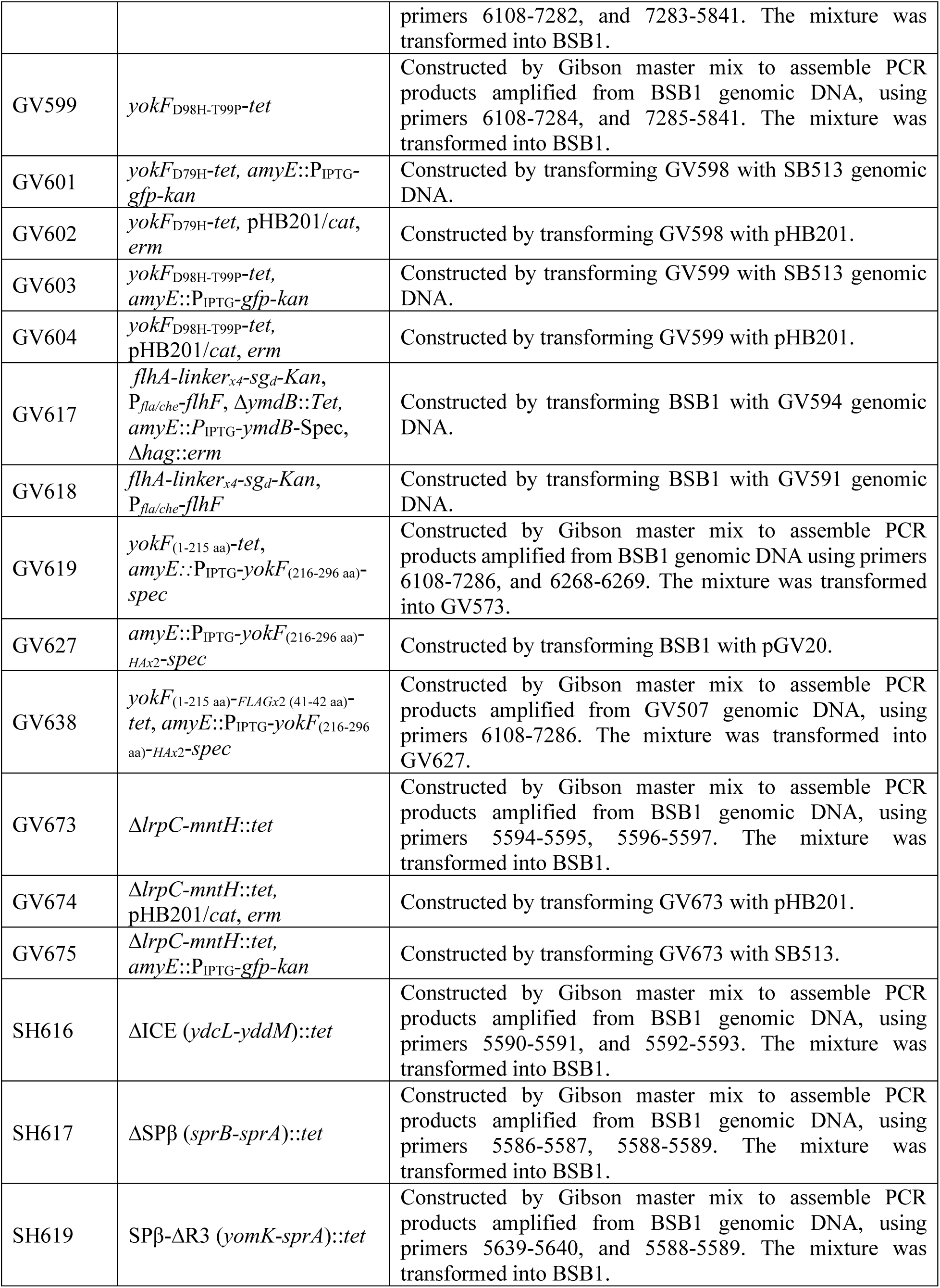
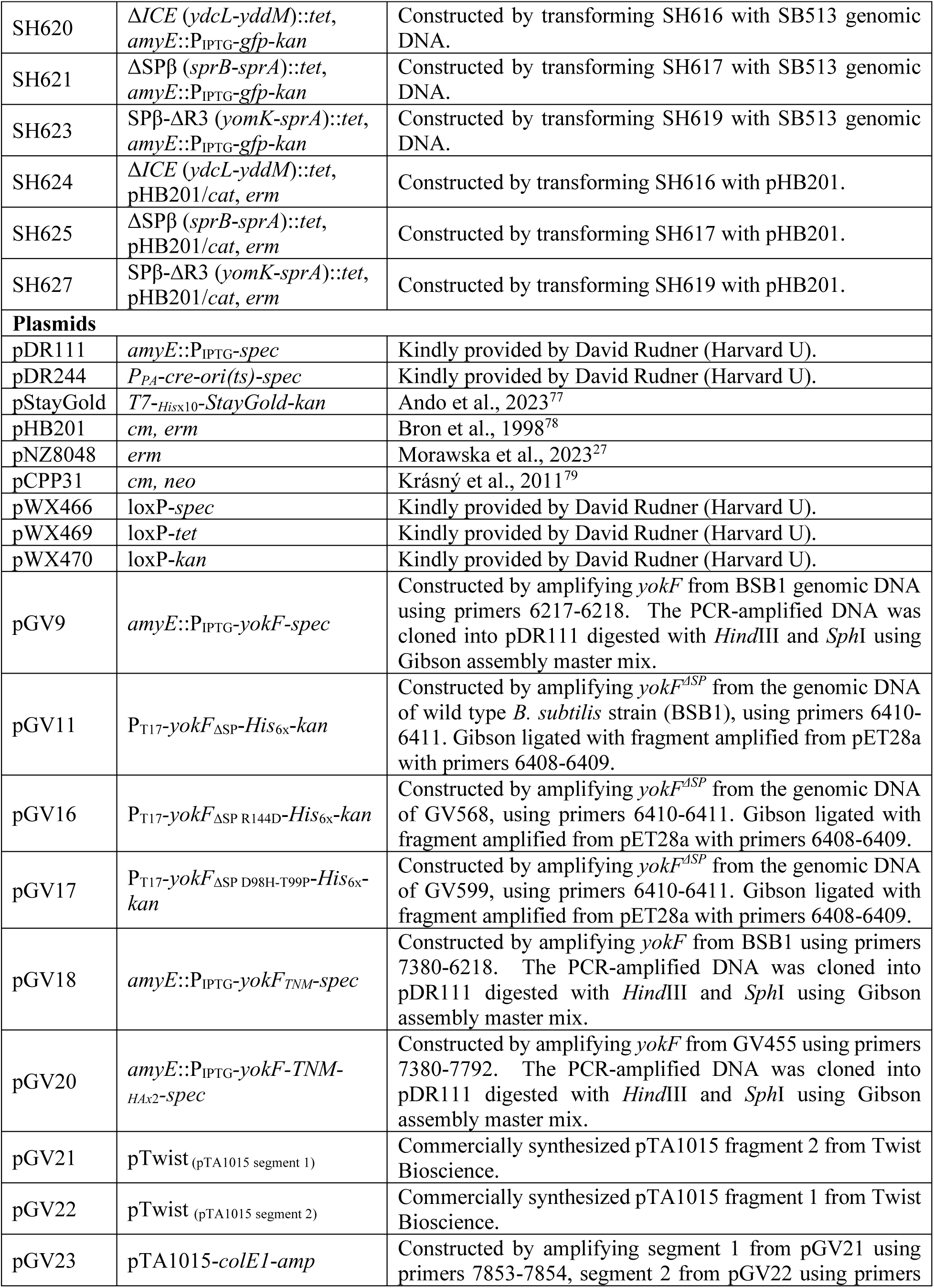
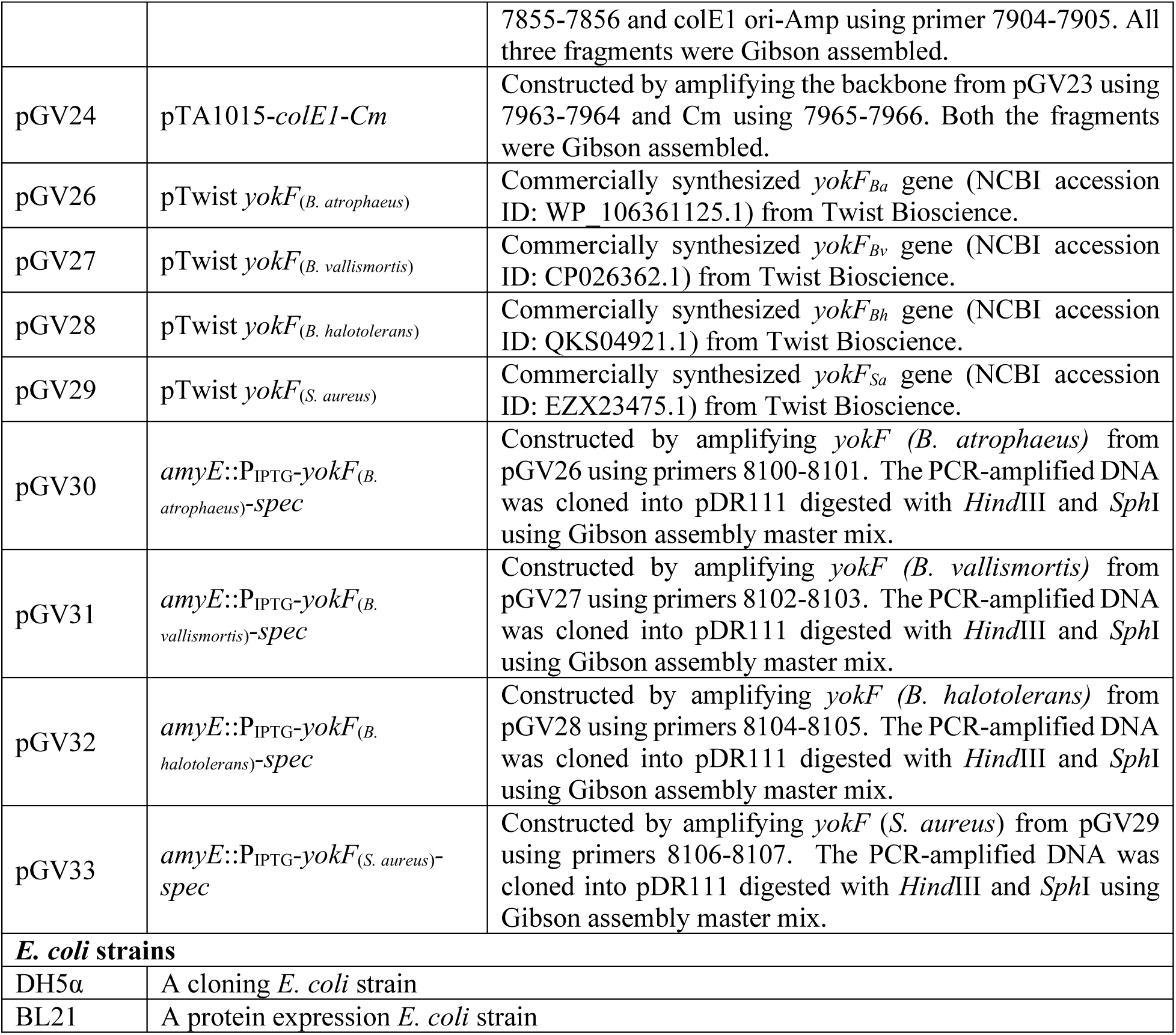
List of strains and plasmids used in this study.

**Table S5.**
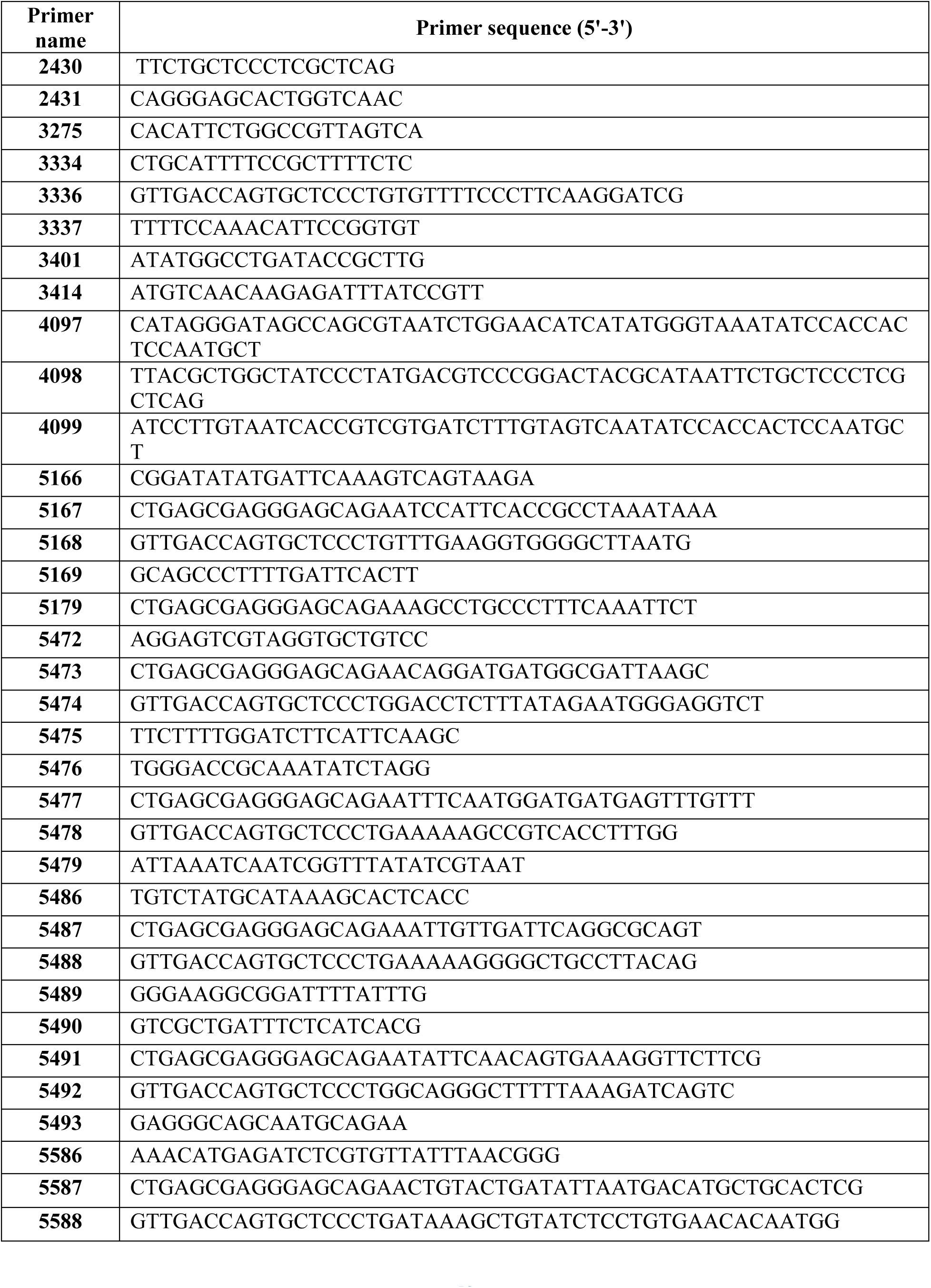
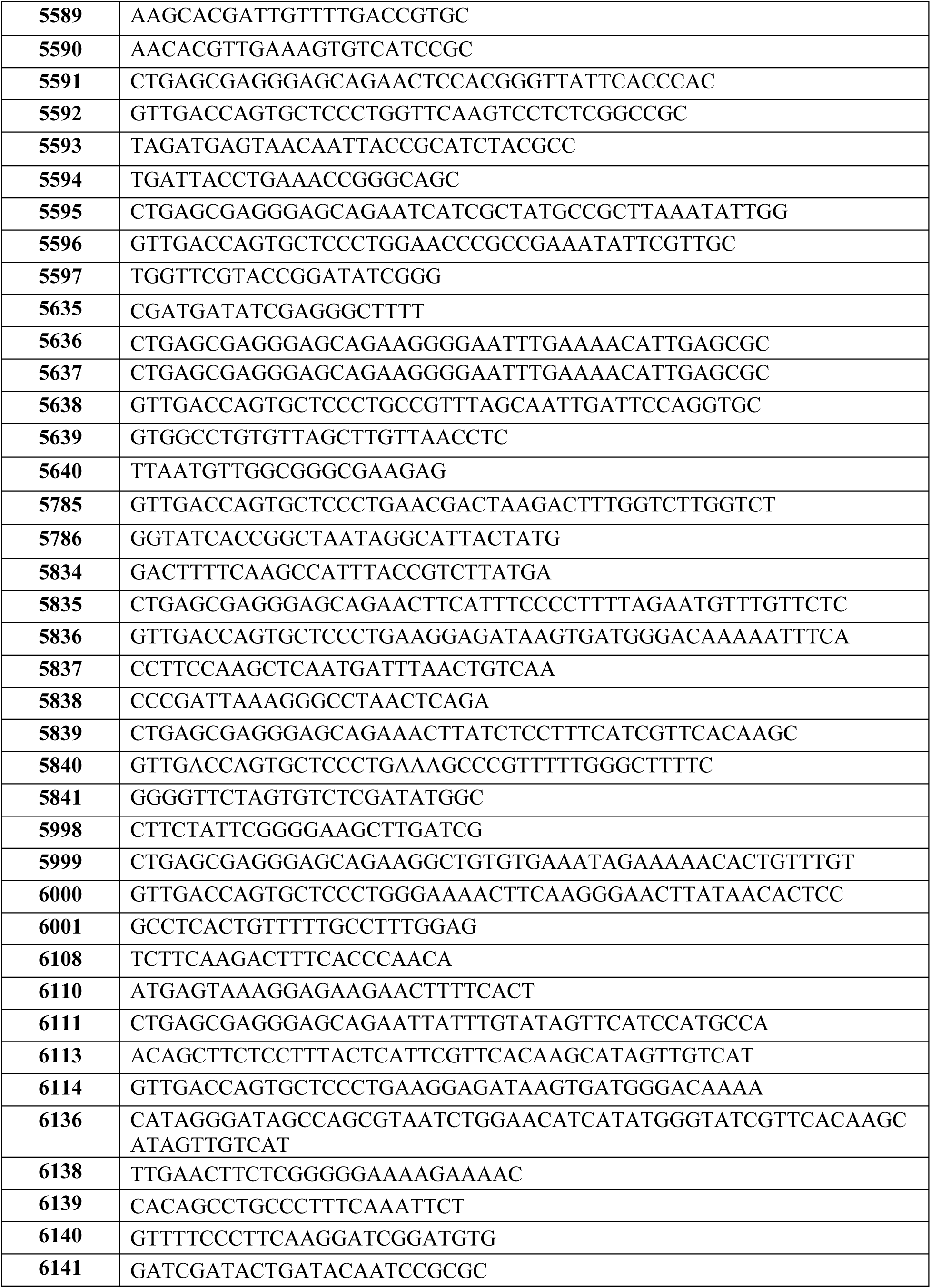
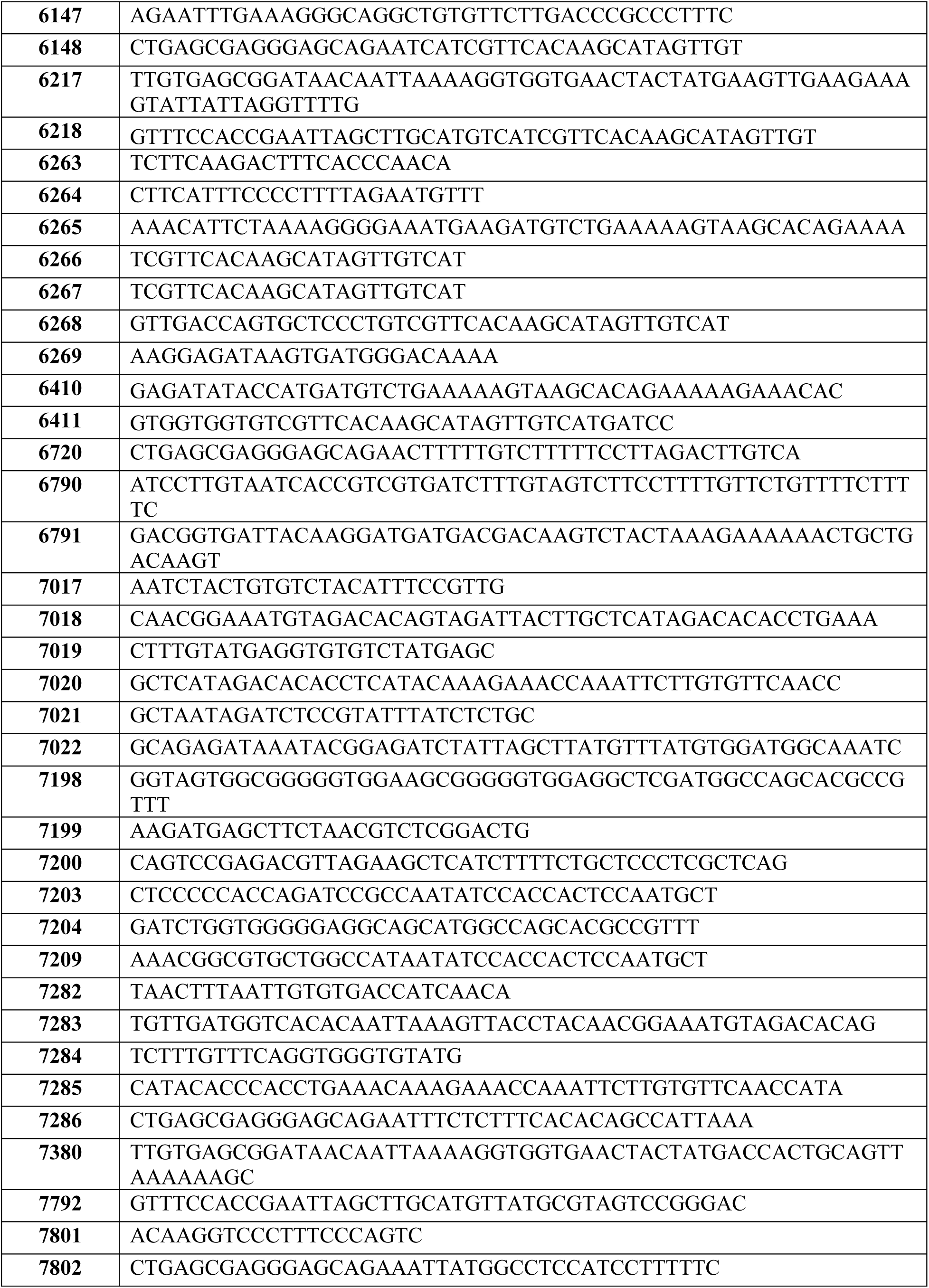
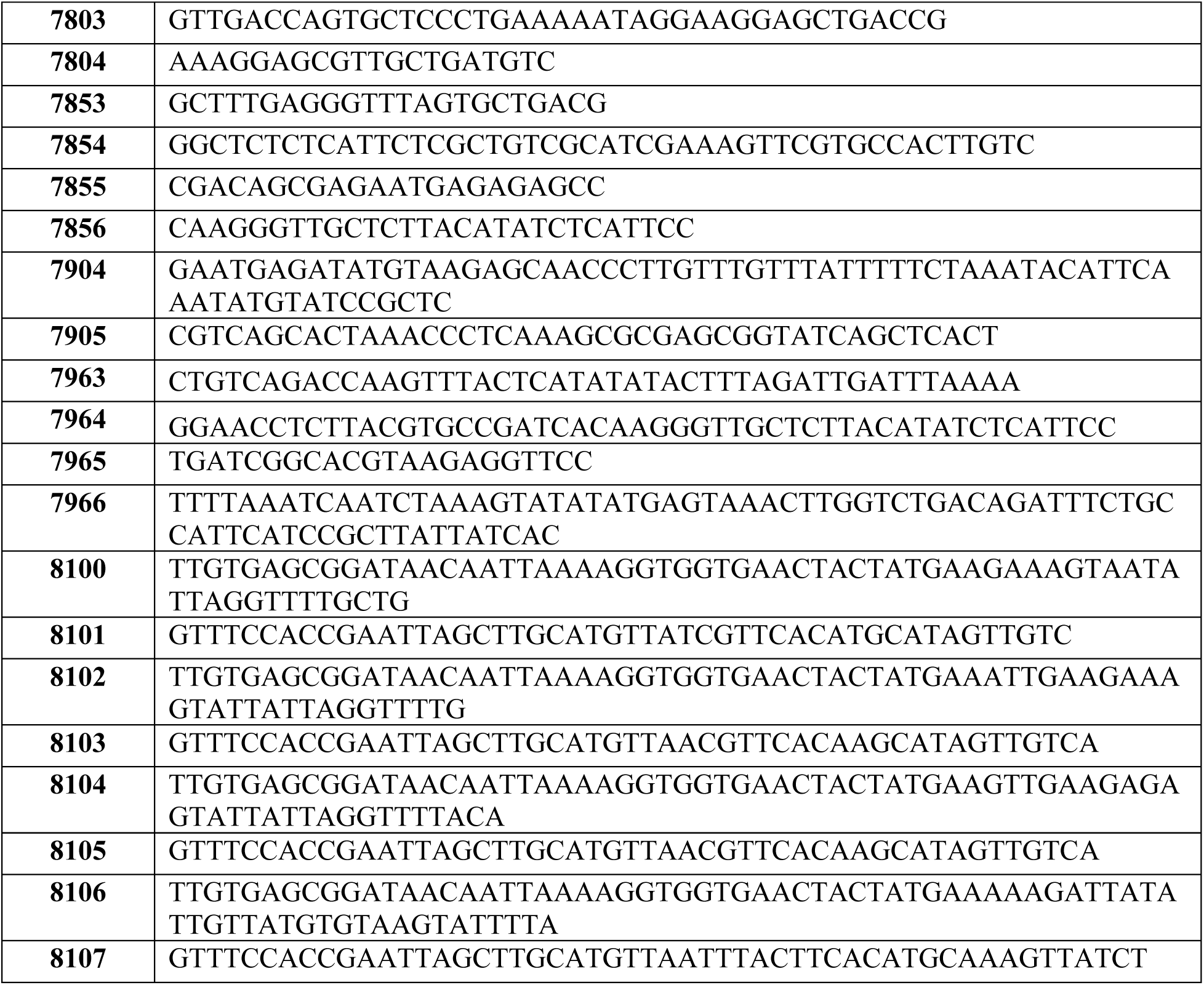
List of primers used in this study.

